# Virus-Receptor Interactions of Glycosylated SARS-CoV-2 Spike and Human ACE2 Receptor

**DOI:** 10.1101/2020.06.25.172403

**Authors:** Peng Zhao, Jeremy L. Praissman, Oliver C. Grant, Yongfei Cai, Tianshu Xiao, Katelyn E. Rosenbalm, Kazuhiro Aoki, Benjamin P. Kellman, Robert Bridger, Dan H. Barouch, Melinda A. Brindley, Nathan E. Lewis, Michael Tiemeyer, Bing Chen, Robert J. Woods, Lance Wells

**Author notes:** Authors contributed equally (Peng Zhao, Jeremy L. Praissman, Oliver C. Grant). To whom correspondence should be addressed: Robert J. Woods, and/or Lance Wells.

## Abstract

The current COVID-19 pandemic is caused by the SARS-CoV-2 betacoronavirus, which utilizes its highly glycosylated trimeric Spike protein to bind to the cell surface receptor ACE2 glycoprotein and facilitate host cell entry. We utilized glycomics-informed glycoproteomics to characterize site-specific microheterogeneity of glycosylation for a recombinant trimer Spike mimetic immunogen and for a soluble version of human ACE2. We combined this information with bioinformatic analyses of natural variants and with existing 3D-structures of both glycoproteins to generate molecular dynamics simulations of each glycoprotein alone and interacting with one another. Our results highlight roles for glycans in sterically masking polypeptide epitopes and directly modulating Spike-ACE2 interactions. Furthermore, our results illustrate the impact of viral evolution and divergence on Spike glycosylation, as well as the influence of natural variants on ACE2 receptor glycosylation that, taken together, can facilitate immunogen design to achieve antibody neutralization and inform therapeutic strategies to inhibit viral infection.

## INTRODUCTION

The SARS-CoV-2 coronavirus, a positive-sense single-stranded RNA virus, is responsible for the severe acute respiratory syndrome referred to as COVID-19 that was first reported in China in December of 2019 (1). In approximately six months, this betacoronavirus has spread globally with more than 14 million people testing positive worldwide resulting in greater than 600,000 deaths as of July 20^th^, 2020 (https://coronavirus.jhu.edu/map.html). The SARS-CoV-2 coronavirus is highly similar (nearly 80% identical at the genomic level) to SARS-CoV-1, which was responsible for the severe acute respiratory syndrome outbreak that began in 2002 (2,3). Furthermore, human SARS-CoV-2 at the whole genome level is >95% identical to a bat coronavirus (RaTG13), the natural reservoir host for multiple coronaviruses (1,4,5). Given the rapid appearance and spread of this virus, there is no current validated vaccine or SARS-CoV-2-specific targeting therapy clinically approved although statins, heparin, and steroids look promising for lowering fatality rates and antivirals likely reduce the duration of symptomatic disease presentation (6-12).

SARS-CoV-2, like SARS-CoV-1, utilizes the host angiotensin converting enzyme II (ACE2) for binding and entry into host cells (13,14). Like many viruses, SARS-CoV-2 utilizes a Spike glycoprotein trimer for recognition and binding to the host cell entry receptor and for membrane fusion (15). Given the importance of viral Spike proteins for targeting and entry into host cells along with their location on the viral surface, Spike proteins are often used as immunogens for vaccines to generate neutralizing antibodies and frequently targeted for inhibition by small molecules that might block host receptor binding and/or membrane fusion (15,16). In similar fashion, wildtype or catalytically-impaired ACE2 has also been investigated as a potential therapeutic biologic that might interfere with the infection cycle of ACE2 targeting coronaviruses (17,18). Thus, a detailed understanding of SARS-CoV-2 Spike binding to ACE2 is critical for elucidating mechanisms of viral binding and entry, as well as for undertaking the rational design of effective therapeutics.

The SARS-CoV-2 Spike glycoprotein consists of two subunits, a receptor binding subunit (S1) and a membrane fusion subunit (S2) (1,2). The Spike glycoprotein assembles into stable homotrimers that together possess 66 canonical sequons for N-linked glycosylation (N-X-S/T, where X is any amino acid except P) as well as a number of potential O-linked glycosylation sites (19,20). Interestingly, coronaviruses virions bud into the lumen of the endoplasmic reticulum-Golgi intermediate compartment, ERGIC, raising unanswered questions regarding the precise mechanisms by which viral surface glycoproteins are processed as they traverse the secretory pathway (21,22). While this and similar studies (19,23) analyze recombinant proteins, a previous study on SARS-CoV-1 suggest that glycosylation of the Spike can be impacted by this intracellular budding and remains to be investigated in SARS-CoV-2 (24). Nonetheless, it has been proposed that this virus, and others, acquires a glycan coat sufficient and similar enough to endogenous host protein glycosylation that it serves as a glycan shield, facilitating immune evasion by masking non-self viral peptides with self-glycans (15,20-22). In parallel with their potential masking functions, glycan-dependent epitopes can elicit specific, even neutralizing, antibody responses, as has been described for HIV-1 ((15,25-29), https://www.biorxiv.org/content/10.1101/2020.06.30.178897v1). Thus, understanding the glycosylation of the viral Spike trimer is fundamental for the development of efficacious vaccines, neutralizing antibodies, and therapeutic inhibitors of infection.

ACE2 is an integral membrane metalloproteinase that regulates the renin-angiotensin system (30). Both SARS-CoV-1 and SARS-CoV-2 have co-opted ACE2 to function as the receptor by which these viruses attach and fuse with host cells (13,14). ACE2 is cleavable by ADAM proteases at the cell surface (31), resulting in the shedding of a soluble ectodomain which can be detected in apical secretions of various epithelial layers (gastric, airway, etc.) and in serum (32). The N-terminal extracellular domain of ACE2 contains 6 canonical sequons for N-linked glycosylation and several potential O-linked sites. Several nonsynonymous single-nucleotide polymorphisms (SNPs) in the ACE2 gene have been identified in the human population and could potentially alter ACE2 glycosylation and/or affinity of the receptor for the viral Spike protein (33).

Given that glycosylation can affect the half-life of circulating glycoproteins in addition to modulating the affinity of their interactions with receptors and immune/inflammatory signaling pathways (34,35), understanding the impact of glycosylation of ACE2 with respect to its binding of SARS-CoV-2 Spike glycoprotein is of high importance. The proposed use of soluble extracellular domains of ACE2 as decoy, competitive inhibitors for SARS-CoV-2 infection emphasizes the critical need for understanding the glycosylation profile of ACE2 so that optimally active biologics can be produced (17,18).

To accomplish the task of characterizing site-specific glycosylation of the trimer Spike of SARS-CoV-2 and the host receptor ACE2, we began by expressing and purifying a stabilized, soluble trimer Spike glycoprotein mimetic immunogen (that we define here and forward as S, (36)) and a soluble version of the ACE2 glycoprotein from a human cell line. We utilized multiple mass spectrometry-based approaches, including glycomic and glycoproteomic approaches, to determine occupancy and site-specific heterogeneity of N-linked glycans. Occupancy (i.e. the percent of any given residue being modified by a glycan) is an important consideration when developing neutralizing antibodies against a glycan-dependent epitope. We also identified sites of O-linked glycosylation and the heterogeneity of the O-linked glycans on S and ACE2. We leveraged this rich dataset, along with existing 3D-structures of both glycoproteins, to generate static and molecular dynamics models of S alone, and in complex with the glycosylated, soluble ACE 2 receptor. By combining bioinformatic characterization of viral evolution and variants of the Spike and ACE2 with molecular dynamics simulations of the glycosylated Spike-ACE2 interaction, we identified important roles for glycans in multiple processes, including receptor-viral binding and glycan-shielding of the Spike. Our rich characterization of the recombinant, glycosylated Spike trimer mimetic immunogen of SARS-CoV-2 in complex with the soluble human ACE2 receptor provides a detailed platform for guiding rational vaccine, antibody, and inhibitor design.

## RESULTS

### Expression, Purification, and Characterization of SARS-CoV-2 Spike Glycoprotein Trimer and Soluble Human ACE2

A trimer-stabilized, soluble variant of the SARS-CoV-2 Spike protein (S) that contains 22 canonical N-linked glycosylation sequons per protomer and a soluble version of human ACE2 that contains 6, lacking the most C-terminal 7th, canonical N-linked glycosylation sequons (**Fig. 1A**) were purified from the media of transfected HEK293 cells and the quaternary structure confirmed by negative EM staining for the S trimer (**Fig. 1B**) and purity examined by SDS-PAGE Coomassie G-250 stained gels for both (**Fig. 1C**). In addition, proteolytic digestions followed by proteomic analyses confirmed that the proteins were highly purified (**Supplemental Table, Tab 12**). Finally, the N-terminus of both the mature S and the soluble mature ACE2 were empirically determined via proteolytic digestions and LC-MS/MS analyses. These results confirmed that both the secreted, mature forms of S protein and ACE2 begin with an N-terminal glutamine that has undergone condensation to form pyroglutamine at residue 14 and 18, respectively (**Figs. 1D and S1**). The N-terminal peptide observed for S also contains a glycan at Asn-0017 (**Fig. 1D**) and mass spectrometry analysis of non-reducing proteolytic digestions confirmed that Cys-0015 of S is in a disulfide linkage with Cys-0136 (**Fig. S2, Supplemental Table, Tab 2**). Given that SignalP (37) predicts signal sequence cleavage between Cys-0015 and Val-0016 but we observed cleavage between Ser-0013 and Gln-0014, we examined the possibility that an in-frame upstream Methionine to the proposed start Methionine (**Fig. 1A**) might be used to initiate translation (**Fig. S3**). If one examines the predicted signal sequence cleavage using the in-frame Met that is encoded 9 amino acids upstream, SignalP now predicts cleavage between the Ser and Gln that we observed in our studies (**Fig. S3**). To examine whether this impacted S expression, we expressed constructs that contained or did not contain the upstream 27 nucleotides in a pseudovirus (VSV) system expressing SARS-CoV2 S (**Fig. S4**) and in our HEK293 system (data not shown). Both expression systems produced a similar amount of S regardless of which expression construct was utilized (**Fig. S4**). Thus, while the translation initiation start site has still not been fully defined, allowing for earlier translation in expression construct design did not have a significant impact on the generation of S.

**Figure 1.**
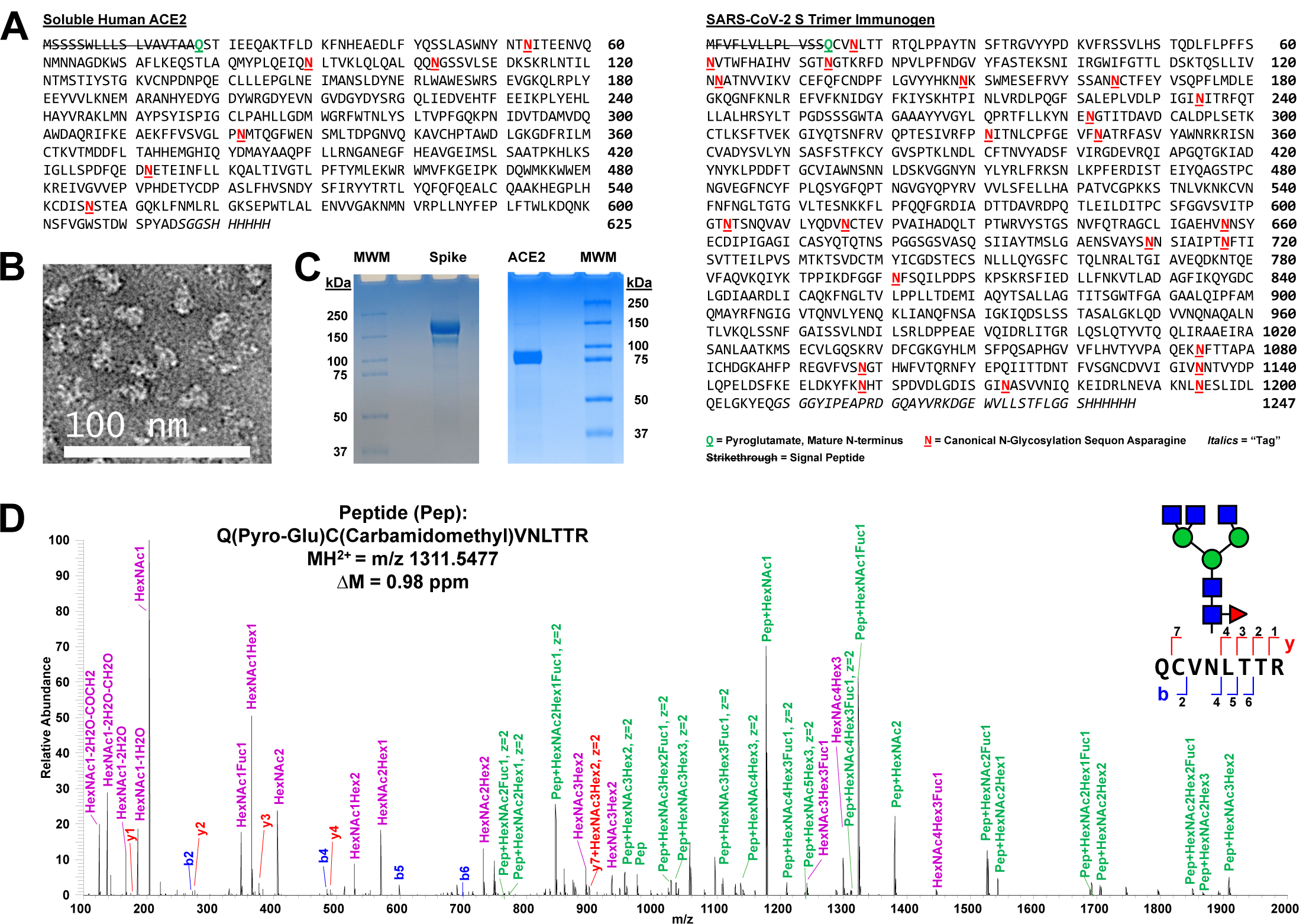
Expression and Characterization of SARS-CoV-2 Spike Glycoprotein Trimer Immunogen and Soluble Human ACE2. A) Sequences of SARS-CoV-2 S immunogen and soluble human ACE2. The N-terminal pyroglutamines for both mature protein monomers are bolded, underlined, and shown in green. The canonical N-linked glycosylation sequons are bolded, underlined, and shown in red. Negative stain electron microscopy of the purified trimer (B) and Coomassie G-250 stained reducing SDS-PAGE gels (C) confirmed purity of the SARS-CoV-2 S protein trimer and of the soluble human ACE2. MWM = molecular weight markers. D) A representative Step-HCD fragmentation spectrum from mass spectrometry analysis of a tryptic digest of S annotated manually based on search results from pGlyco 2.2. This spectrum defines the N-terminus of the mature protein monomer as (pyro-)glutamine 0014. A representative N-glycan consistent with this annotation and our glycomics data (Fig. 2) is overlaid using the Symbol Nomenclature For Glycans (SNFG) code. This complex glycan occurs at N0017. Note, that as expected, the cysteine is carbamidomethylated and the mass accuracy of the assigned peptide is 0.98 ppm. On the sequence of the N-terminal peptide and in the spectrum, the assigned b (blue) and y (red) ions are shown. In the spectrum, purple highlights glycan oxonium ions and green marks intact peptide fragment ions with various partial glycan sequences still attached. Note that the green-labeled ions allow for limited topology to be extracted including defining that the fucose is on the core and not the antennae of the glycopeptide.

### Glycomics Informed Glycoproteomics Reveals Site-Specific Microheterogeneity of SARS-CoV-2 S Glycosylation

We utilized multiple approaches to examine glycosylation of the SARS-CoV-2 S trimer. First, the portfolio of glycans linked to SARS-CoV-2 S trimer immunogen was analyzed following their release from the polypeptide backbone. N-glycans were released from protein by treatment with PNGase F and O-glycans were subsequently released by beta-elimination. Following permethylation to enhance detection sensitivity and structural characterization, released glycans were analyzed by multi-stage mass spectrometry (MS^n^) (38,39). Mass spectra were processed by GRITS Toolbox and the resulting annotations were validated manually (40). Glycan assignments were grouped by type and by additional structural features for relative quantification of profile characteristics (**Fig. 2A, Supplemental Table, Tab 3**). This analysis quantified 49 N-glycans and revealed that 55% of the total glycan abundance was of the complex type, 17% was of the hybrid type, and 28% was high-mannose. Among the complex and hybrid N-glycans, we observed a high degree of core fucosylation and significant abundance of bisected and LacDiNAc structures. We also observed sulfated N-linked glycans using negative mode MS^n^ analyses (**Supplemental Table, Tab 13**) though signal intensity was too low in positive ion mode (at least 10-fold lower than any of the non-sulfated glycans) for accurate quantification. In addition, we detected 15 O-glycans released from the S trimer (**Fig. S5, Supplemental Table, Tab 4**).

**Figure 2.**
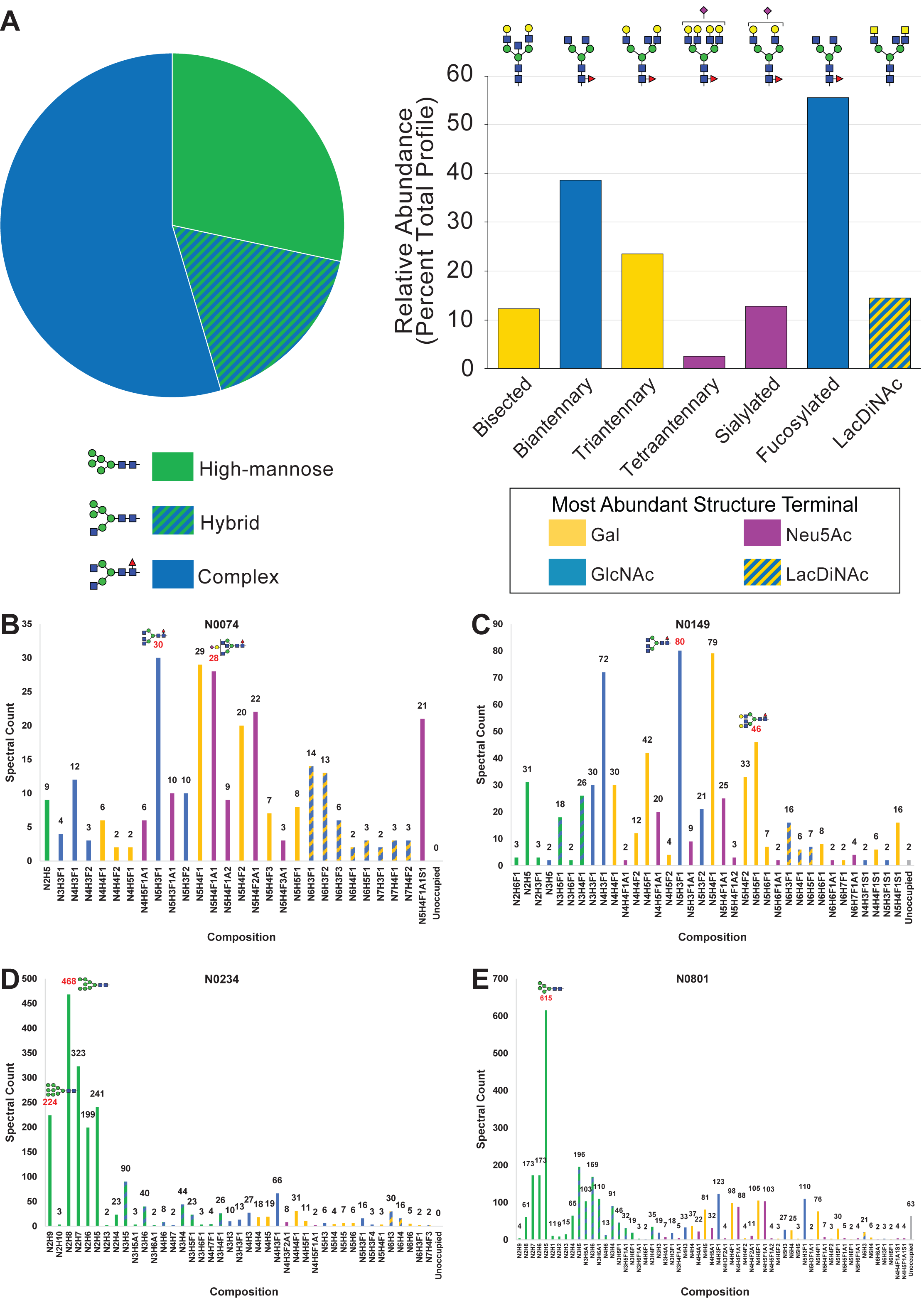
Glycomics Informed Glycoproteomics Reveals Substantial Site-Specific Microheterogeneity of N-linked Glycosylation on SARS-CoV-2 S. A) Glycans released from SARS-CoV-2 S protein trimer immunogen were permethylated and analyzed by MSn. Structures were assigned, grouped by type and structural features, and prevalence was determined based on ion current. The pie chart shows basic division by broad N-glycan type. The bar graph provides additional detail about the glycans detected. The most abundant structure with a unique categorization by glycomics for each N-glycan type in the pie chart, or above each feature category in the bar graph, is indicated. B – E) Glycopeptides were prepared from SARS-CoV-2 S protein trimer immunogen using multiple combinations of proteases, analyzed by LC-MSn, and the resulting data was searched using several different software packages. Four representative sites of N-linked glycosylation with specific features of interest were chosen and are presented here. N0074 (B) and N0149 (C) are shown that occur in variable insert regions of S compared to SARS-CoV and other related coronaviruses, and there are emerging variants of SARS-CoV-2 that disrupt these two sites of glycosylation in S. N0234 (D) contains the most high-mannose N-linked glycans. N0801 (D) is an example of glycosylation in the S2 region of the immunogen and displays a high degree of hybrid glycosylation compared to other sites. The abundance of each composition is graphed in terms of assigned spectral counts. Representative glycans (as determined by glycomics analysis) for several abundant compositions are shown in SNFG format. The abbreviations used here and throughout the manuscript are N for HexNAc, H for Hexose, F for Fucose, A for Neu5Ac, and S for Sulfation. Note that the graphs for the other 18 sites and other graphs grouping the microheterogeneity observed by other properties are presented in Supplemental Information.

To determine occupancy of N-linked glycans at each site, we employed a sequential deglycoslyation approach using Endoglycosidase H and PNGase F in the presence of ^18^O-H_2_O following tryptic digestion of S (28,41). Following LC-MS/MS analyses, the resulting data confirmed that 19 of the canonical sequons had occupancies greater than 95% (**Supplemental Table, Tab 5**). One canonical sequence, N0149, had insufficient spectral counts for quantification by this method but subsequent analyses described below suggested high occupancy. The 2 most C-terminal N-linked sites, N1173 and N1194, had reduced occupancy, 52% and 82% respectively.

Reduced occupancy at these sites may reflect hindered en-bloc transfer by the oligosaccharyltransferase (OST) due to primary amino acid sequences at or near the N-linked sequon. Alternatively, this may reflect these two sites being post-translationally modified after release of the protein by the ribosome by a less efficient STT3B-containing OST, either due to activity or initial folding of the polypeptide, as opposed to co-translationally modified by the STT3A-containing OST (42). None of the non-canoncial sequons (3 N-X-C sites and 4 N-G-L/I/V sites, (43)) showed significant occupancy (>5%) except for N0501 that showed moderate (19%) conversion to ^18^O-Asp that could be due to deamidation that is facilitated by glycine at the +1 position (**Supplemental Table, Tab 5**, (44)). Further analysis of this site (see below) by direct glycopeptide analyses allowed us to determine that N0501 undergoes deamidation but is not glycosylated. Thus, all, and only the, 22 canonical sequences for N-linked glycosylation (N-X-S/T) are utilized with only N1173 and N1194 demonstrating occupancies below 95%.

Next, we applied 3 different proteolytic digestion strategies to the SARS-CoV-2 S immunogen to maximize glycopeptide coverage by subsequent LC-MS/MS analyses. Extended gradient nanoflow reverse-phase LC-MS/MS was carried out on a ThermoFisher Lumos(tm) Tribrid(tm) instrument using Step-HCD fragmentation on each of the samples (see STAR methods for details, (25,26,28,41,45)). Following data analyses using pGlyco 2.2.2 (46), Byonic (47), and manual validation of glycan compositions against our released glycomics findings (**Fig. 2A, Supplemental Table, Tab 3 and 13**), we were able to determine the microheterogeneity at each of the 22 canonical sites (**Fig. 2B-2E, Supplemental Table, Tab 6**). Notably, none of the non-canonical consensus sequences, including N0501, displayed any quantifiable glycans. The N-glycosites N0074 (**Fig. 2B**) and N0149 (**Fig. 2C**) are highly processed and display a typical mammalian N-glycan profile. N0149 is, however, modified with several hybrid N-glycan structures while N0074 is not. N0234 (**Fig. 2D**) and N0801 (**Fig. 2E**) have N-glycan profiles more similar to those found on other viruses such as HIV (15) that are dominated by high-mannose structures. N0234 (**Fig. 2D**) displays an abundance of Man7 - Man9 high-mannose structures suggesting stalled processing by early acting ER and cis-Golgi mannosidases. In contrast, N0801 (**Fig. 2E**) is processed more efficiently to Man5 high-mannose and hybrid structures suggesting that access to the glycan at this site by MGAT1 and *α*-Mannosidase II is hindered. In general, for all 22 sites (**Fig. 2B-2E, Supplemental Table, Tab 6**), we observed under processing of complex glycan antennae (i.e. under-galactosylation and under-sialylation) and a high degree of core fucosylation in agreement with released glycan analyses (**Fig. 2A, Supplemental Table, Tab 3**). We also observed a small percent of sulfated N-linked glycans at several sites (**Supplemental Table, Tab 6 and 8**). Based on the assignments and the spectral counts for each topology, we were able to determine the percent of total N-linked glycan types (high-mannose, hybrid, or complex) present at each site (**Figure 3, Supplemental Table, Tab 7**). Notably, 3 of the sites (N0234, N0709, and N0717) displayed more than 50% high-mannose glycans while 11 other sites (N0017, N0074, N0149, N0165, N0282, N0331, N0657, N1134, N1158, N1173, and N1194) were more than 90% complex when occupied. The other 8 sites were distributed between these 2 extremes. Notably, only 1 site (N0717 at 45%), which also had greater than 50% high-mannose (55%), had greater than 33% hybrid structures. To further evaluate the heterogeneity, we grouped all the topologies into the 20 classes recently described by the Crispin laboratory with adding 2 categories (sulfated and unoccupied) that we refer to here as the Oxford classification (**Supplemental Table, Tab 8**, (19)). Among other features observed, this classification allowed us to observe that while most sites with high mannose structures were dominated by the Man5GlcNAc2 structure, N0234 and N0717 were dominated by the higher Man structures of Man8GlcNAc2 and Man7GlcNAc2, respectively (**Fig. S7, Supplemental Table, Tab 8**). Limited processing at N0234 is in agreement with a recent report suggesting that high mannose structures at this site help to stabilize the receptor-binding domain of S (www.biorxiv.org/content/10.1101/2020.06.11.146522v1). Furthermore, applying the Oxford classifications to our dataset clearly demonstrates that the 3 most C-terminal sites (N1158, N1173, and N1194), dominated by complex type glycans, were more often further processed (i.e. multiple antennae) and elaborated (i.e. galactosylation and sialylation) than other sites (**Supplemental Table, Tab 8**).

**Figure 3.**
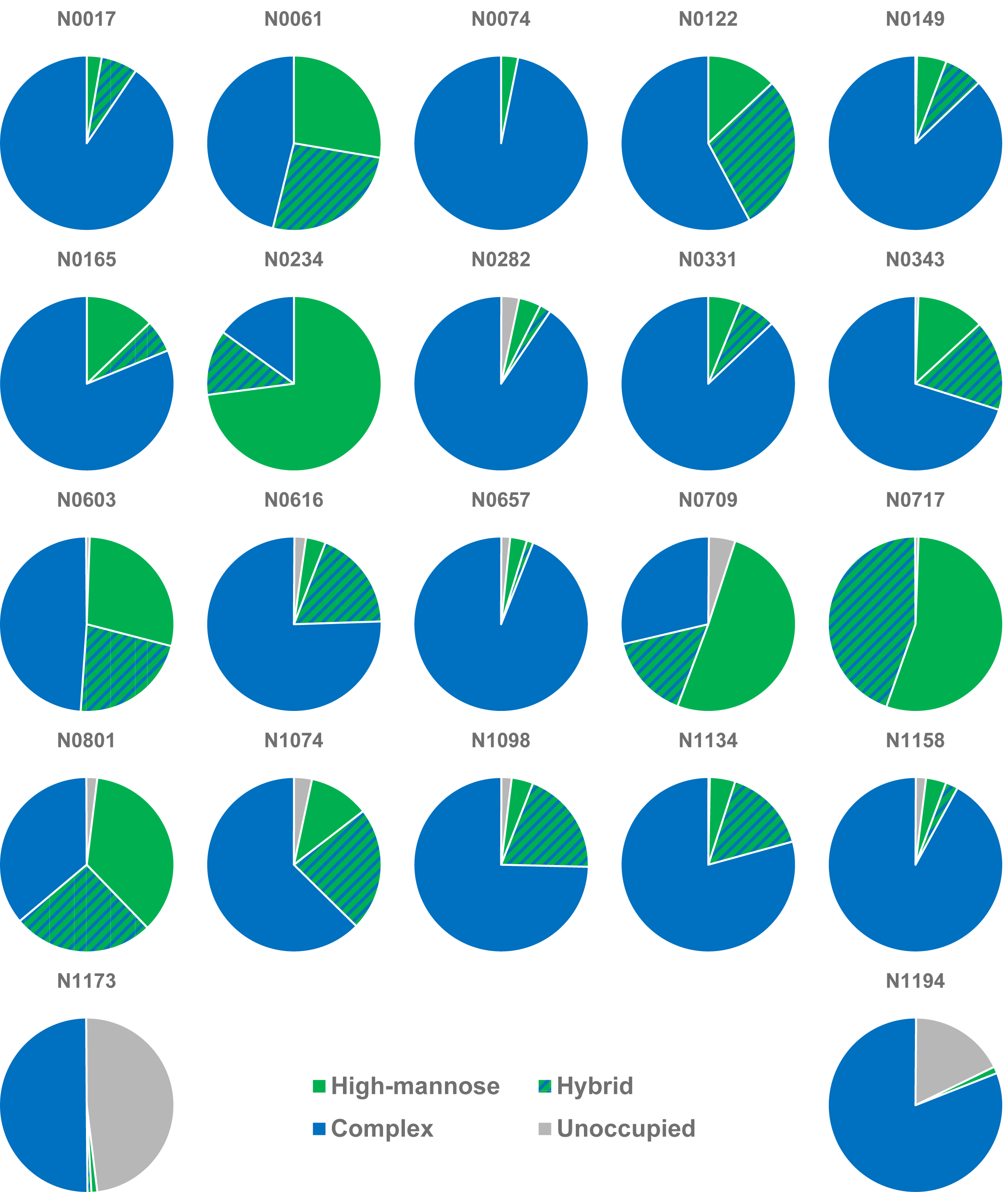
SARS-CoV-2 S Immunogen N-glycan Sites are Predominantly Modified by Complex N-glycans. N-glycan topologies were assigned to all 22 sites of the S protomer and the spectral counts for each of the 3 types of N-glycans (high-mannose, hybrid, and complex) as well as the unoccupied peptide spectral match counts at each site were summed and visualized as pie charts. Note that only N1173 and N1194 show an appreciable amount of the unoccupied amino acid.

We also analyzed our generated mass spectrometry data for the presence of O-linked glycans based on our glycomic findings (**Fig. S5, Supplemental Table, Tab 4**) and a recent manuscript suggesting significant levels of O-glycosylation of S1 and S2 when expressed independently (23). We were able to confirm sites of O-glycan modification with microheterogeneity observed for the vast majority of these sites (**Supplemental Table, Tab 9**). However, occupancy at each site, determined by spectral counts, was observed to be very low (below 4%) except for Thr0323 that had a modestly higher but still low 11% occupancy (**Fig. S6, Supplemental Table, Tab 10**).

### 3D Structural Modeling of Glycosylated SARS-CoV-2 Trimer Immunogen Enables Predictions of Epitope Accessibility and Other Key Features

A 3D structure of the S trimer was generated using a homology model of the S trimer described previously (based on PDB code 6VSB, (48)). Onto this 3D structure, we installed explicitly defined glycans at each glycosylated sequon based on one of three separate sets of criteria, thereby generating three different glycoform models for comparison that we denote as “Abundance,” “Oxford Class,” and “Processed” models (**see Methods and Supplemental Table, Tab 1**). These criteria were chosen in order to generate glycoform models that represent reasonable expectations for glycosylation microheterogeneity and integrate cross-validating glycomic and glycoproteomic characterization of S and ACE2.

The three glycoform models were subjected to multiple all atom MD simulations with explicit water. Information from analyses of these structures is presented in **Figure 4A** along with the sequence of the SARS-CoV-2 S protomer. We also determined variants in S that are emerging in the virus that have been sequenced to date (**Supplemental Table, Tab 11**). The inter-residue distances were measured between the most *α*-carbon-distal atoms of the N-glycan sites and Spike glycoprotein population variant sites in 3D space (**Figure 4B**). Notable from this analysis, there are several variants that don’t ablate the N-linked sequon, but that are sufficiently close in 3-dimensional space to N-glycosites, such as D138H, H655Y, S939F, and L1203F, to warrant further investigation.

**Figure 4.**
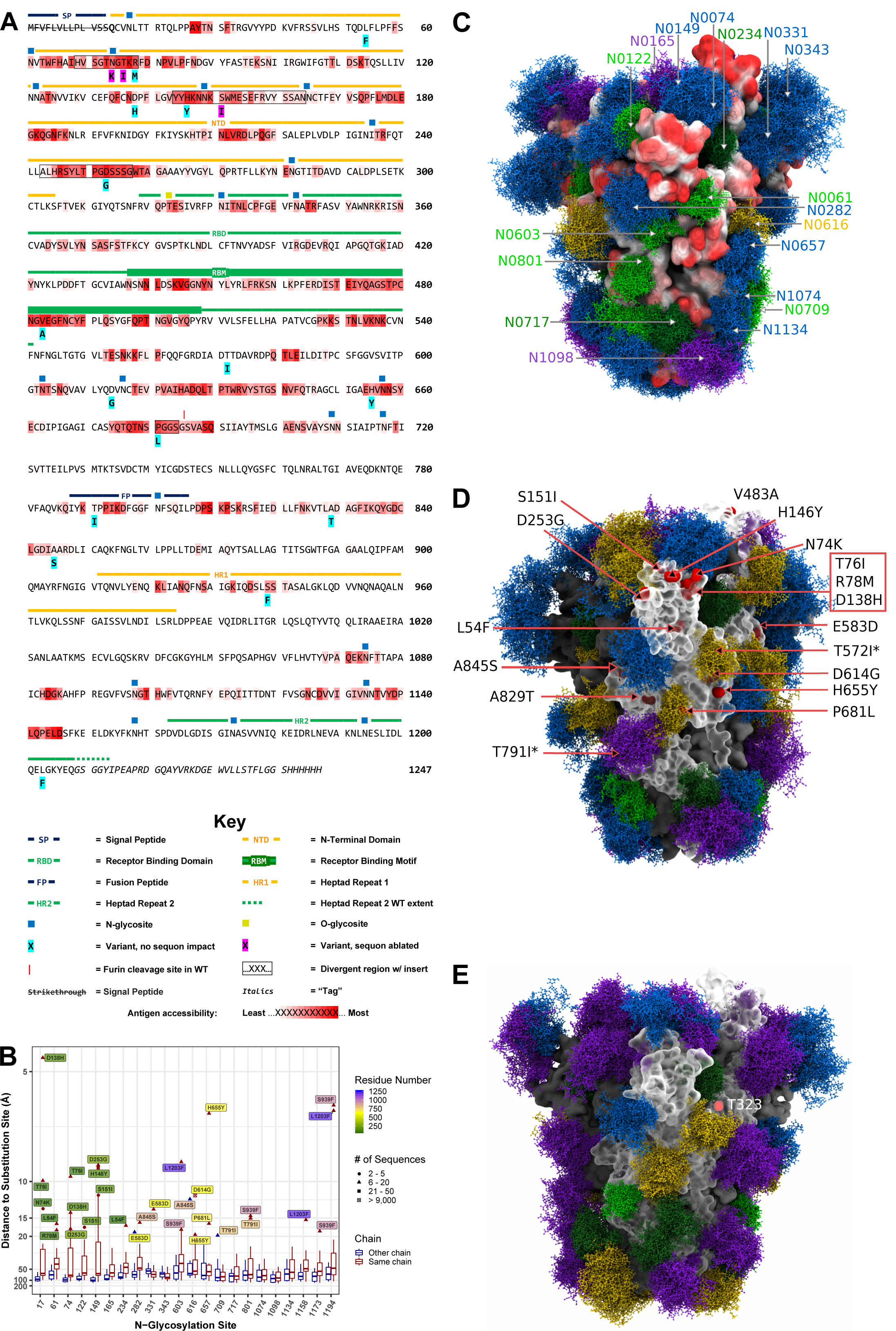
3D Structural Modeling of Glycosylated SARS-CoV-2 Spike Trimer Immunogen Reveals Predictions for Antigen Accessibility and Other Key Features. Results from glycomics and glycoproteomics experiments were combined with results from bioinformatics analyses and used to model several versions of glycosylated SARS-CoV-2 S trimer immunogen. A) Sequence of the SARS-CoV-2 S immunogen displaying computed antigen accessibility and other information. Antigen accessibility is indicated by red shading across the amino acid sequence. B) Emerging variants confirmed by independent sequencing experiments were analyzed based on the 3D structure of SARS-CoV-2 S to generate a proximity chart to the determined N-linked glycosylation sites. C) SARS-CoV-2 S trimer immunogen model from MD simulation displaying abundance glycoforms and antigen accessibility shaded in red for most accessible, white for partial, and black for inaccessible (see **Supplemental movie A**). D) SARS-CoV-2 S trimer immunogen model from MD simulation displaying oxford class glycoforms and sequence variants. * indicates not visible while the box represents 3 amino acid variants that are clustered together in 3D space. E) SARS-CoV-2 S trimer immunogen model from MD simulation displaying processed glycoforms plus shading of Thr-323 that has O-glycoslyation at low stoichiometry in yellow.

The percentage of simulation time that each S protein residue is accessible to a probe that approximates the size of an antibody variable domain was calculated for a model of the S trimer using the Abundance glycoforms (**Supplemental Table, Tab 1**, (49)). The predicted antibody accessibility is visualized across the sequence, as well as mapped onto the 3D surface, via color shading **(Figure 4A, 4C, Supplemental Table, Tab 13, and Supplemental Movie A)**. Additionally, the Oxford Class glycoforms model (**Supplemental Table, Tab 1)**, which is arguably the most encompassing means for representing glycan microheterogeneity since it captures abundant structural topologies (**Supplemental Table, Tab 8**), is shown with the sequence variant information (Figure 4D, **Supplemental Table, Tab 11**). A substantial number of these variants occur (directly by comparison to **Figure 4A** or visually by comparison to **Figure 4C**) in regions of high calculated epitope accessibility (e.g. N74K, T76I, R78M, D138H, H146Y, S151I, D253G, V483A, etc., **Supplemental Table, Tab 14**) suggesting potential selective pressure to avoid host immune response. Also, it is interesting to note that 3 of the emerging variants would eliminate N-linked sequons in S; N74K and T76I would eliminate N-glycosylation of N74 (found in the insert variable region 1 of CoV-2 S compared to CoV-1 S), and S151I eliminates N-glycosylation of N149 (found in the insert variable region 2) (**Fig. 4A, S7, Supplemental Table, Tab 11)**. Lastly, the SARS-CoV-2 S Processed glycoform model is shown (**Supplemental Table, Tab 1**), along with marking amino acid T0323 that has a modest (11% occupancy, **Fig. S6, Supplemental Table, Tab 10**) amount of O-glycosylation to represent the most heavily glycosylated form of S (**Figure 4E**).

### Glycomics Informed Glycoproteomics Reveals Complex N-linked Glycosylation of ACE2

We also analyzed ACE2 glycosylation utilizing the same glycomic and glycoproteomic approaches described for S protein. Glycomic analyses of released N-linked glycans (**Fig. 5A, Supplemental Table, Tab 3**) revealed that the majority of glycans on ACE2 are complex with limited high-mannose and hybrid glycans and we were unable to detect sulfated N-linked glycans. Glycoproteomic analyses revealed that occupancy was high (>75%) at all 6 sites and significant microheterogeneity dominated by complex N-glycans was observed for each site (**Fig. 5B-5G, Supplemental Table, Tabs 5-8**). We also observed, consistent with the O-glycomics (**Fig. S5, Supplemental Table, Tab 4**), that Ser 155 and several S/T residues at the C-terminus of ACE2 outside of the peptidase domain were O-glycosylated but stoichiometry was extremely low (less than 2%, **Supplemental Table, Tab 9 and 10**).

**Figure 5:**
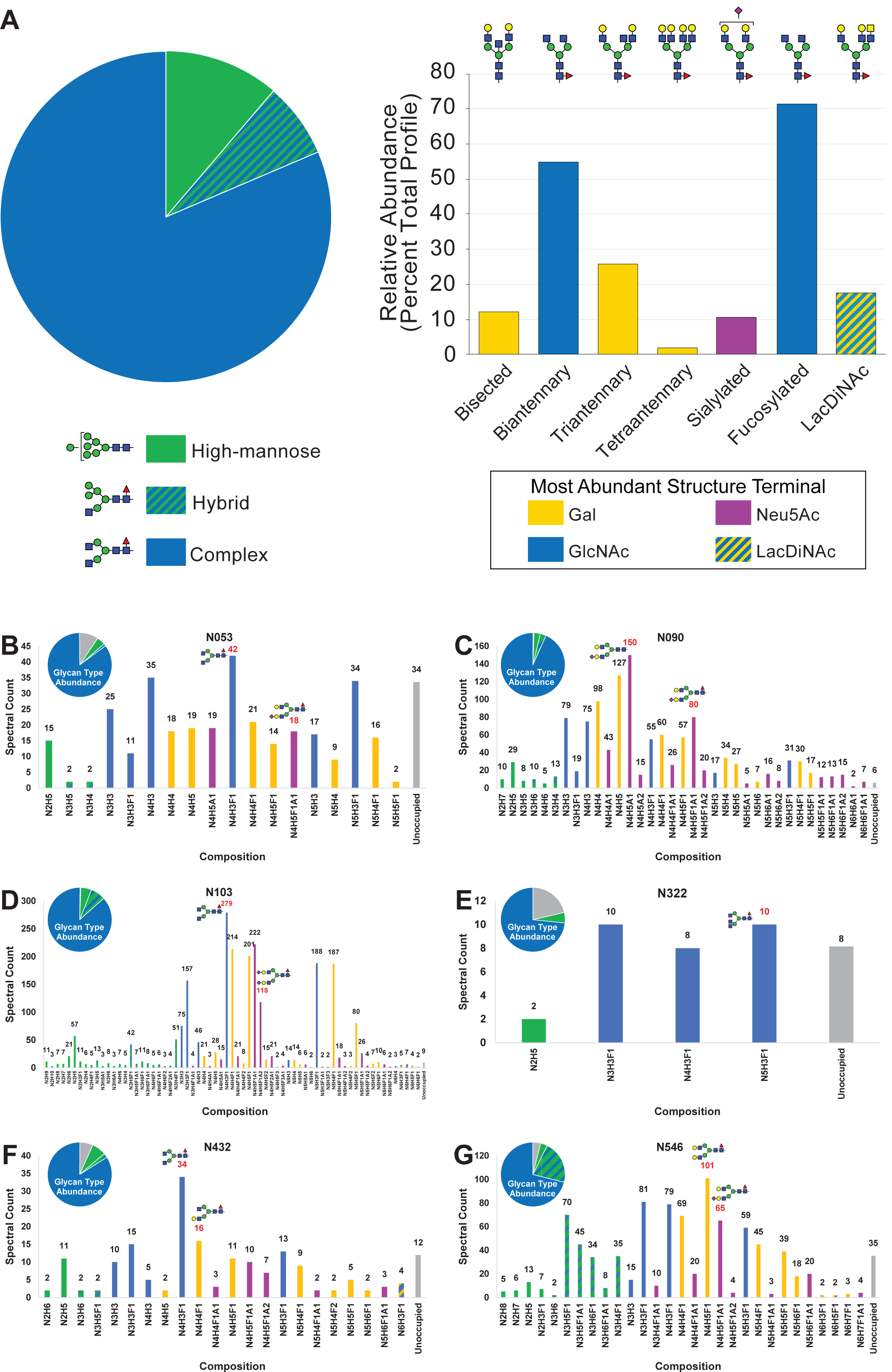
Glycomics Informed Glycoproteomics of Soluble Human ACE2 Reveals High Occupancy, Complex N-linked Glycosylation. A) Glycans released from soluble, purified ACE2 were permethylated and analyzed by MSn. Structures were assigned, grouped by type and structural features, and prevalence was determined based on ion current. The pie chart shows basic division by broad N-glycan type. The bar graph provides additional detail about the glycans detected. The most abundant structure with a unique categorization by glycomics for each N-glycan type in the pie chart, or above each feature category in the bar graph, is indicated. B – G) Glycopeptides were prepared from soluble human ACE2 using multiple combinations of proteases, analyzed by LC-MSn, and the resulting data was searched using several different software packages. All six sites of N-linked glycosylation are presented here. Displayed in the bar graphs are the individual compositions observed graphed in terms of assigned spectral counts. Representative glycans (as determined by glycomics analysis) for several abundant compositions are shown in SNFG format. The abbreviations used here and throughout the manuscript are N for HexNAc, H for Hexose, F for Fucose, and A for Neu5Ac. The pie chart (analogous to Figure 3 for SARS-CoV-2 S) for each site is displayed in the upper corner of each panel. B) N053. C) N090. D) N103. E) N322. F) N432. G) N546, a site that does not exist in 3 in 10,000 people.

### 3D Structural Modeling of Glycosylated, Soluble, ACE2 Highlighting Glycosylation and Variants

We integrated our glycomics, glycoproteomics, and population variant analyses results with a 3D model of Ace 2 (based on PDB code 6M0J (50), see methods for details) to generate two versions of the soluble glycosylated ACE2 for visualization and molecular dynamics simulations. We visualized the ACE2 glycoprotein with the Abundance glycoform model simulated at each site as well as highlighting the naturally occurring variants observed in the human population (**Fig. 6A, Supplemental Movie B, Supplemental Table, Tab 11**). Note, that the Abundance glycoform model and the Oxford Class glycoform model for ACE2 are identical (**Supplemental Table, Tabs 1 and 8**). Notably, one site of N-linked glycosylation (N546) is predicted to not be present in 3 out of 10,000 humans based on naturally occurring variation in the human population (**Supplemental Table, Tab 11**). We also modeled ACE2 using the Processed glycoform model (**Fig. 6B**). In both models, the interaction domain with S is defined (**Fig. 6A-B, Supplemental Movie B**).

**Figure 6:**
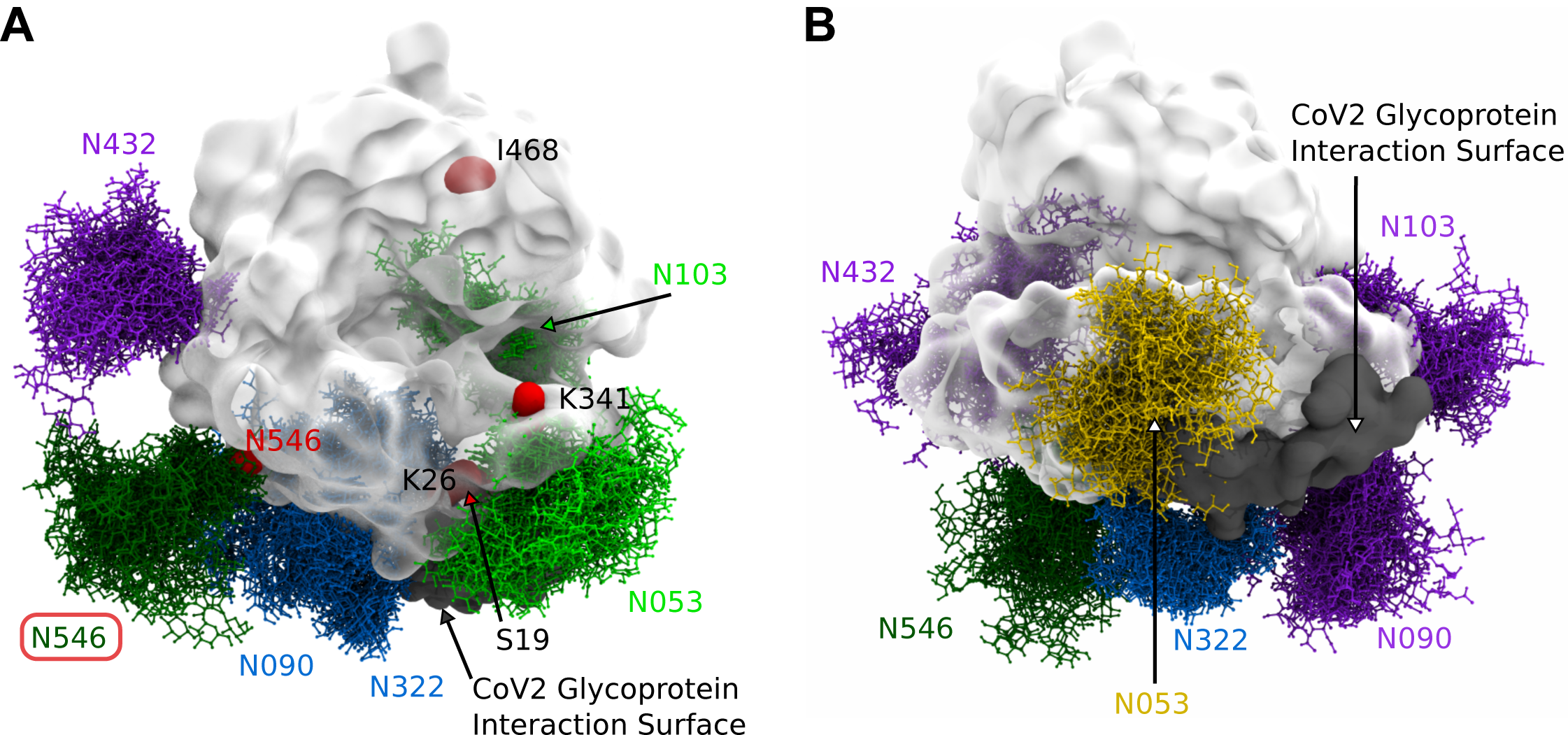
3D Structural Modeling of Glycosylated Soluble Human ACE2. Results from glycomics and glycoproteomics experiments were combined with results from bioinformatics analyses and used to model several versions of glycosylated soluble human ACE2. A) Soluble human ACE2 model from MD simulations displaying abundance glycoforms, interaction surface with S, and sequence variants. N546 variant is boxed that would remove N-linked glycosylation at that site (see **Supplemental movie B**). B) Soluble human ACE2 model from MD simulations displaying processed glycoforms and interaction surface with S.

### Molecular Dynamics Simulation of the Glycosylated Trimer Spike of SARS-CoV-2 in Complex with Glycosylated, Soluble, Human Ace 2 Reveals Protein and Glycan Interactions

Molecular dynamics simulations were performed to examine the co-complex (generated from a crystal structure of the ACE2-RBD co-complex, PDB code 6M0J, (50)) of glycosylated S with glycosylated ACE2 with the 3 different glycoforms models (Abundance, Oxford Class, and Processed, **Supplemental Table, Tab 1, Supplemental Simulations 1-3**). Information from these analyses is laid out along the primary structure (sequence) of the SARS-CoV-2 S protomer and ACE2 highlighting regions of glycan-protein interaction observed in the MD simulations (**Supplemental Table, Tab 14, Supplemental Simulations 1-3**). Interestingly, two glycans on ACE2 (at N090 and N322), that are highlighted in **Figure 7A** and shown in a more close-up view in **Figure 7B**, are predicted to form interactions with the S protein (**Supplemental Table, Tab 15**). The N322 glycan interaction with the S trimer is outside of the receptor binding domain, and the interaction is observed across multiple simulations and throughout each simulation (**Fig. 7A-B, Supplemental Simulations 1-3**). The ACE2 glycan at N090 is close enough to the S trimer surface to repeatedly form interactions, however the glycan arms interact with multiple regions of the surface over the course of the simulations, reflecting the relatively high degree of glycan dynamics (**Fig. 7A-B, Supplemental Movie C**). Inter-molecule glycan-glycan interactions are also observed repeatedly between the glycan at N546 of ACE2 and those in the S protein at residues N0074 and N0165 (**Fig. 7D, Supplemental Table, Tab 16**). Finally, a full view of the ACE2-S complex with Oxford class glycoforms on both proteins illustrates the extensive glycosylation at the interface of the complex (**Fig. 7C, Supplemental Movie D**).

**Figure 7:**
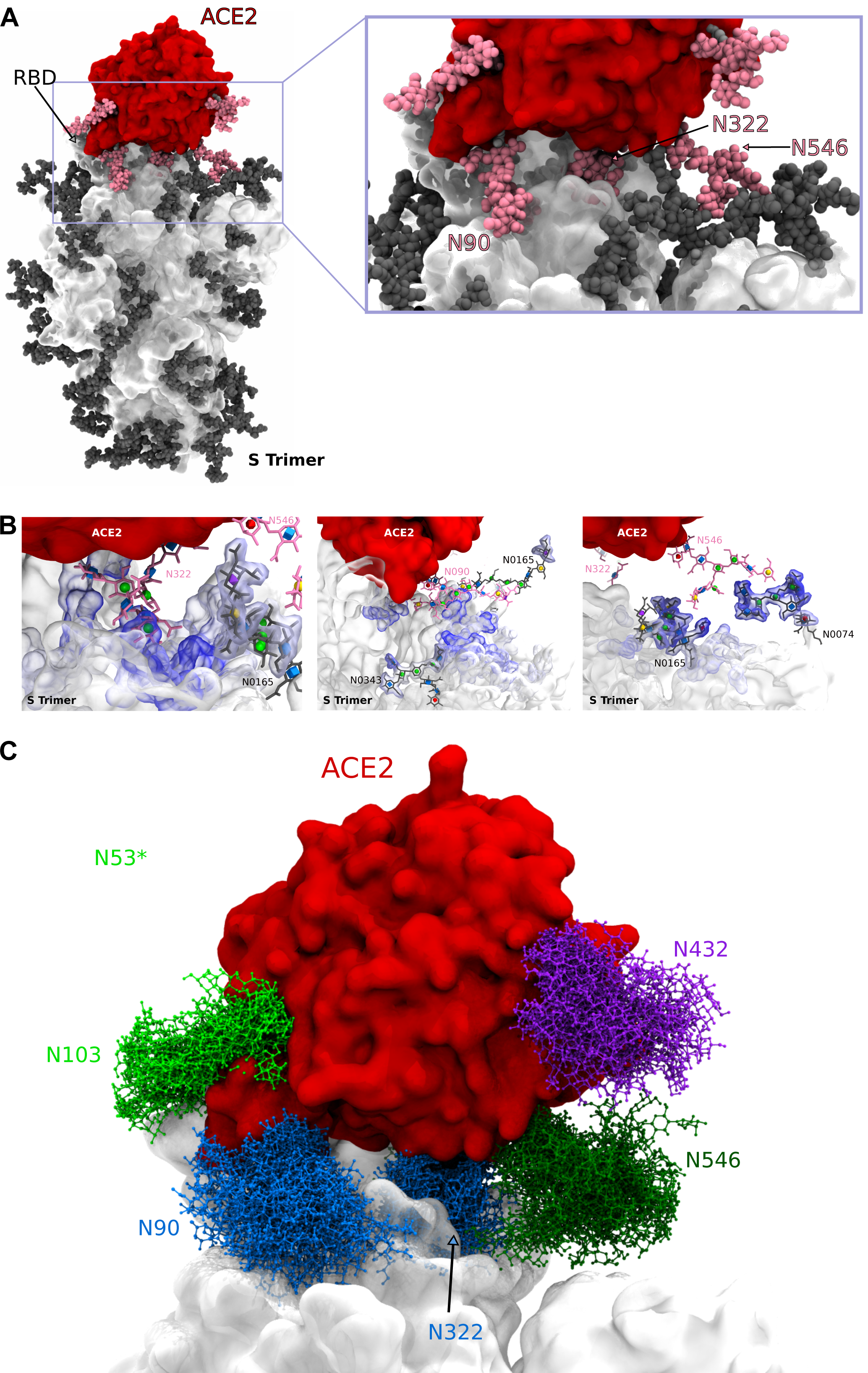
Interactions of Glycosylated Soluble Human ACE2 and Glycosylated SARS-CoV-2 S Trimer Immunogen Revealed By 3D-Structural Modeling and Molecular Dynamics Simulations. A) Molecular dynamics simulation of glycosylated soluble human ACE2 and glycosylated SARS-CoV-2 S trimer immunogen interaction (see **Supplemental simulations 1-3**). ACE2 (top) is colored red with glycans in pink while S is colored white with glycans in dark grey. Highlighted are ACE2 glycans that interacts with S that are zoomed in on to the right. B) Zoom in of ACE2-S interface highlighting ACE2 glycan interactions using 3D-SNFG icons (60) with S protein (pink) as well as ACE2-S glycan-glycan interactions. C) Zoom in of dynamics trajectory of glycans at the interface of soluble human ACE2 and S (see **Supplemental movies C and D**).

## DISCUSSION

We have defined the glycomics-informed, site-specific microheterogeneity of 22 sites of N-linked glycosylation per monomer on a SARS-CoV-2 trimer and the 6 sites of N-linked glycosylation on a soluble version of its human ACE2 receptor using a combination of mass spectrometry approaches coupled with evolutionary and variant sequence analyses to provide a detailed understanding of the glycosylation states of these glycoproteins (**Figs. 1-6**). Our results suggest essential roles for glycosylation in mediating receptor binding, antigenic shielding, and potentially the evolution/divergence of these glycoproteins.

The highly glycosylated SARS-CoV-2 Spike protein, unlike several other viral proteins including HIV-1 (15) but in agreement with another recent report (19), presents significantly more processing of N-glycans towards complex glycosylation, suggesting that steric hindrance to processing enzymes is not a major factor at most sites (**Figs. 2-3**). However, the N-glycans still provide considerable shielding of the peptide backbone (**Fig. 4**). Our glycomics-guided glycoproteomic data is in general in strong agreement with the trimer immunogen data recently published by Crispin (19) though we also observed sulfated N-linked glycans, were able to differentiate branching, bisected, and diLacNAc containing structures by glycomics, and observed less occupancy on the 2 most C-terminal N-linked sites using a different approach. Our detection of sulfated N-linked glycans at multiple sites on S is in agreement with a recent manuscript re-analyzing the Crispin data (https://www.biorxiv.org/content/10.1101/2020.05.31.125302v1). Sulfated N-linked glycans could potentially play key roles in immune regulation and receptor binding as in other viruses (51). This result is especially significant in that sulfated N-glycans were not observed when we performed glycomics on ACE2. At each individual site, the glycans we observed on our immunogen appear to be slightly more processed but the overlap between our analysis and the Crispin’s group results (19) at each site in terms of major features are nearly superimposable. This agreement differs substantially when comparing our and Crispin’s data (19) to that of the Azadi group (23) that analyzed S1 and S2 that had been expressed individually. When expressed as 2 separate polypeptides and not purified for trimers, several unoccupied sites of N-linked glycosylation were observed and processing at several sites was significantly different (23) than we and others (19) observed. Although O-glycosylation has recently been reported for individually-expressed S1 and S2 domains of the Spike glycoprotein (23), in trimeric form the level of O-glycosylation is extremely low, with the highest level of occupancy we observed being 11% at T0323 (**Fig. 4E**). The low level of O-linked occupancy we observed is in agreement with Crispin’s analysis of a Spike Trimer immunogen (19) but differs significantly from Azadi’s analyses of individually expressed S1 and S2 (23). Thus, the context in which the Spike protein is expressed and purified before analyses significantly alters the glycosylation of the protomer that is reminiscent of previous studies looking at expression of the HIV-1 envelope Spike (15,52). The soluble ACE2 protein examined here contains 6 highly utilized sites of N-linked glycosylation dominated by complex type N-linked glycans (**Fig. 5**). O-glycans were also present on this glycoprotein but at very low levels of occupancy at all sites (<2%).

Our glycomics-informed glycoproteomics allowed us to assign defined sets of glycans to specific glycosylation sites on 3D-structures of S and ACE2 glycoproteins based on experimental evidence (**Figs. 4, 6**). Similar to almost all glycoproteins, microheterogeneity is evident at most glycosylation sites of S and ACE2; each glycosylation site can be modified with one of several glycan structures, generating site-specific glycosylation portfolios. For modeling purposes, however, explicit structures must be placed at each glycosylation site. In order to capture the impact of microheterogenity on S and ACE2 molecular dynamics we chose to generate glycoforms for modeling that represented reasonable portfolios of glycan types. Using 3 glycoform models for S (Abundance, Oxford Class, and Processed) and 2 models for ACE2 (Abundance, which was equivalent to Oxford Class, and Processed), we generated 3 molecular dynamics simulations of the co-complexes of these 2 glycoproteins (**Fig. 7 and Supplemental Simulations 1-3**). The observed interactions over time allowed us to evaluate glycan-protein contacts between the 2 proteins as well as examine potential glycan-glycan interactions (**Fig. 7**). We observed glycan-mediated interactions between the S trimer and glycans at N090, N322 and N546 of ACE2. Thus, variations in glycan occupancy or processing at these sites, could alter the affinity of the SARS-CoV-2 – ACE2 interaction and modulate infectivity. It is well established that glycosylation states vary depending on tissue and cell type as well as in the case of humans, on age (53), underlying disease (54,55) and ethnicity (56). Thus glycosylation portfolios may in part be responsible for tissue tropism and individual susceptibility to infection. The importance of glycosylation for S binding to ACE2 is even more emphatically demonstrated by the direct glycan-glycan interactions observed (**Fig. 7**) between S glycans (at N0074 and N0165) and an ACE2 receptor glycan (at N546), adding an additional layer of complexity for interpreting the impact of glycosylation on individual susceptibility.

Several emerging variants of the virus appear to be altering N-linked glycosylation occupancy by disrupting N-linked sequons. Interestingly, the 2 N-linked sequons in SARS-CoV-2 S directly impacted by variants, N0074 and N0149, are in divergent insert regions 1 and 2, respectively, of SARS-CoV-2 S compared to SARS-CoV-1 S (**Fig. 4A**). The N0074, in particular, is one of the S glycans that interact directly with ACE2 glycan (at N546, **Fig. 7**), suggesting that glycan-glycan interactions may contribute to the unique infectivity differences between SARS-CoV-2 and SARS-CoV-1. These sequon variants will also be important to examine in terms of glycan shielding that could influence immunogenicity and efficacy of neutralizing antibodies, as well as interactions with the host cell receptor ACE2. Naturally-occurring amino acid-changing SNPs in the ACE2 gene generate a number of variants including 1 variant, with a frequency of 3 in 10,000 humans, that eliminates a site of N-linked glycosylation at N546 (**Fig. 6**). Understanding the impact of ACE2 variants on glycosylation and more importantly on S binding, especially for N546S which impacts the glycan-glycan interaction between S and ACE2 (**Fig. 7**), should be prioritized in light of efforts to develop ACE2 as a potential decoy therapeutic. Intelligent manipulation of ACE2 glycosylation may lead to more potent biologics capable of acting as better competitive inhibitors of S binding. The data presented here, and related similar recent findings (19,57,58), provide a framework to facilitate the production of immunogens, vaccines, antibodies, and inhibitors as well as providing additional information regarding mechanisms by which glycan microheterogeneity is achieved. However, considerable efforts still remain in order to fully understand the role of glycans in SARS-CoV-2 infection and pathogenicity. While HEK-expressed S and ACE2 provide a useful window for understanding human glycosylation of these proteins, glycoproteomic characterization following expression in cell lines of more direct relevance to disease and target tissue is sorely needed. While site occupancy may change depending on presentation and cell type (59), processing of N-linked glycans will almost certainly be altered in a cell-type dependent fashion. Thus, analyses of the Spike trimer extracted from pseudoviruses, virion-like particles, and ultimately from infectious SARS-CoV-2 virions harvested from airway cells or patients will provide the most accurate view of how trimer immunogens reflect the true glycosylation pattern of the virus. Detailed analyses of the impact of emerging variants in S and natural and designed-for-biologics variants of ACE2 on glycosylation and binding properties are important next steps for developing therapeutics. Finally, it will be important to monitor the slow evolution of the virus to determine if existing sites of glycosylation are lost or new sites emerge with selective pressure that might alter the efficacy of vaccines, neutralizing antibodies, and/or inhibitors.

## Supporting information

Supplemental Simulation 1

Supplemental Simulation 2

Supplemental Simulation 3

## ACKNOWLEDGEMENTS

The authors would like to thank Protein Metrics for providing licenses for their software used here and the developers of pGlyco for productive discussions regarding their software. We would also like to thank Galit Alter of the Ragon Institute for facilitating this collaborative effort. This effort was facilitated by the ThermoFisher Scientific appointed Center of Excellence in Glycoproteomics at the Complex Carbohydrate Research Center at the University of Georgia (co-directed by MT and LW). This research is supported in part by R35GM119850 (NEL), NNF10CC1016517 (NEL), R01AI139238 (MAB), R01AI147884-01A1S1 (BC), Massachusetts Consortium on Pathogen Readiness (BC), U01CA207824 (R.J.W), P41GM103390 (R.J.W.), P41GM103490 (MT and LW), U01GM125267 (MT), and R01GM130915 (LW). The content is solely the responsibility of the authors and does not necessarily represent the official views of the National Institutes of Health.

## AUTHOR CONTRIBUTIONS

Conceptualization: M.T., B.C., R.J.W. and L.W.; Methodology, Software, Validation, Formal Analysis, Investigation, Resources, and Data Curation: P.Z., J.L.P., O.C.G., Y.C., T.X., K.E.R., K.A., B.P.K., R.B., D.H.B., M.A.B., N.E.L., M.T., B.C., R.J.W., and L.W.; Writing-Original Draft: P.Z., J.L.P., and L.W.; Writing-Review & Editing: All authors; Visualization: P.Z., J.L.P., O.C.G, Y.C., M.T., B.C., R.J.W., and L.W.; Supervision, Project Administration, and Funding Acquisition: D.H.B., M.A.B., N.E.L., M.T., B.C., R.J.W., and L.W..

## DECLARATION OF INTERESTS

The authors declare no competing interests.

## STAR METHODS

### KEY RESOURCES TABLE

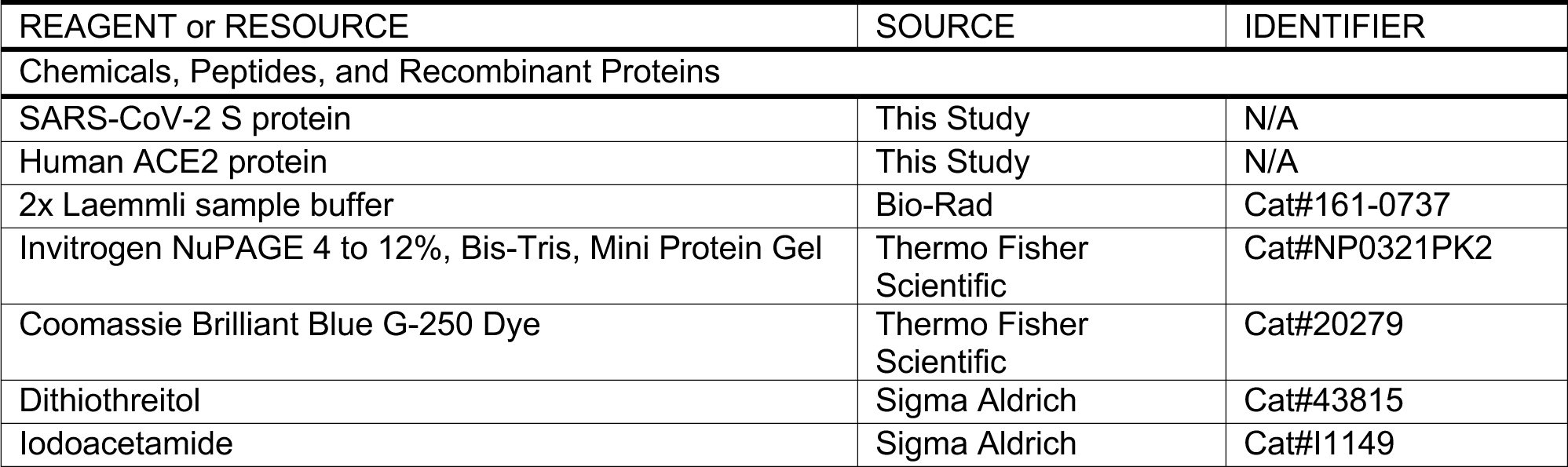
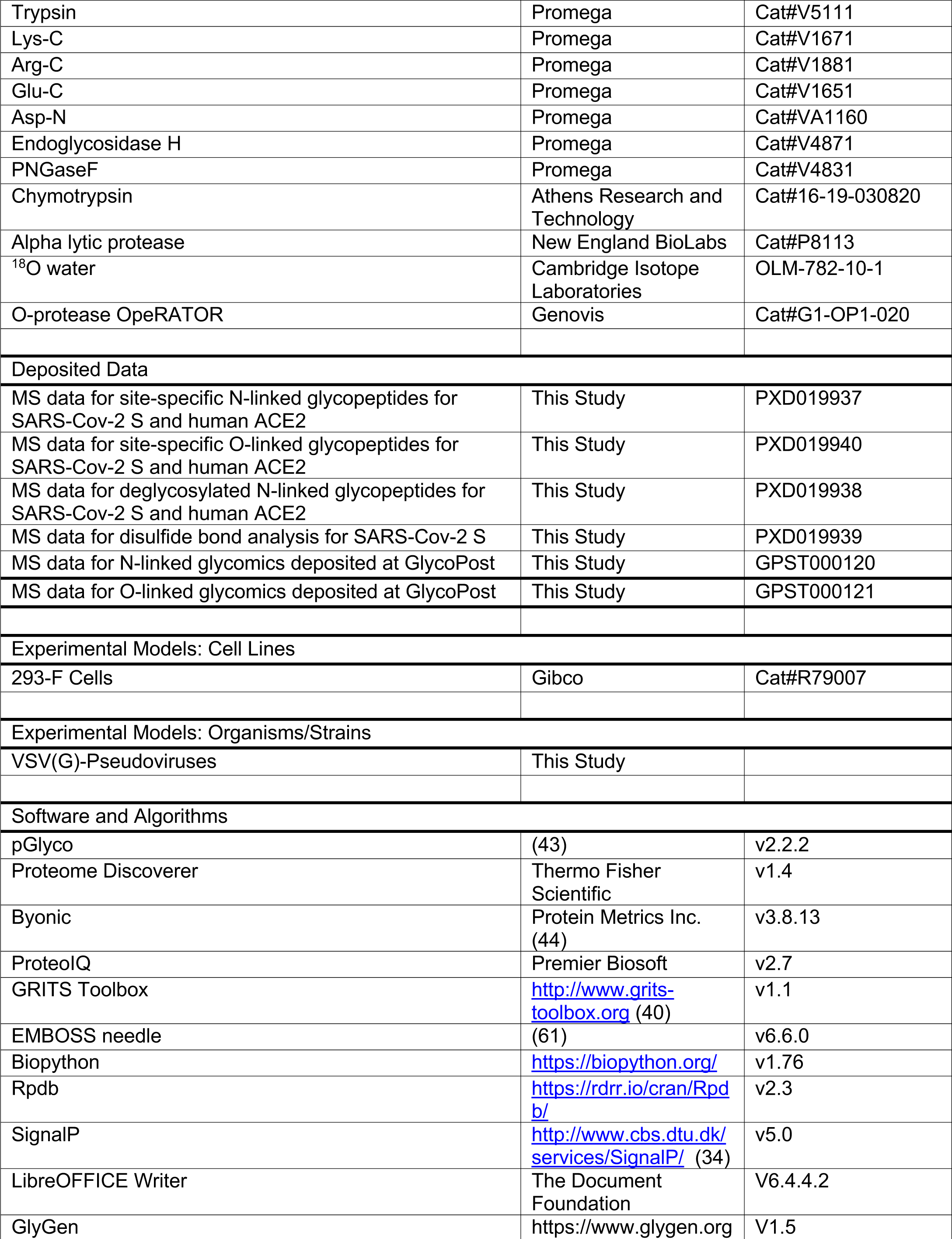
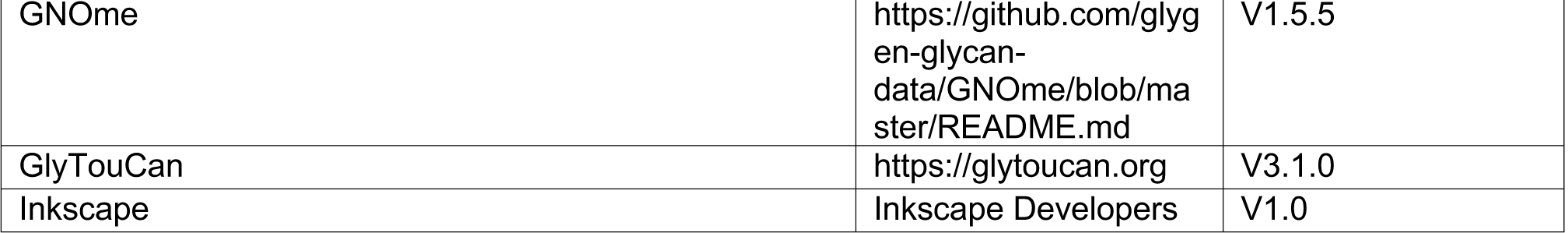

### LEAD CONTACT

Further information and requests for resources and reagents should be directed to and will be fulfilled by the Lead Contact, Peng Zhao and/or Lance Wells.

## METHOD DETAILS

### Expression, Purification, and Characterization of SARS-CoV-2 S and Human ACE2 Proteins

To express a stabilized ectodomain of Spike protein, a synthetic gene encoding residues 1–1208 of SARS-CoV-2 Spike with the furin cleavage site (residues 682–685) replaced by a “GGSG” sequence, proline substitutions at residues 986 and 987, and a foldon trimerization motif followed by a C-terminal 6xHisTag was created and cloned into the mammalian expression vector pCMV-IRES-puro (Codex BioSolutions, Inc, Gaithersburg, MD). The expression construct was transiently transfected in HEK 293F cells using polyethylenimine (Polysciences, Inc, Warrington, PA). Protein was purified from cell supernatants using Ni-NTA resin (Qiagen, Germany), the eluted fractions containing S protein were pooled, concentrated, and further purified by gel filtration chromatography on a Superose 6 column (GE Healthcare). Negative stain electron microscopy (EM) analysis was performed as described (62). Briefly, analysis was performed at room temperature with a magnification of 52,000x and a defocus value of 1.5 µm following low-dose procedures, using a Philips Tecnai F20 electron microscope (Thermo Fisher Scientific) equipped with a Gatan US4000 CCD camera and operated at voltage of 200 kV. The DNA fragment encoding human ACE2 (1-615) with a 6xHis tag at C terminus was synthesized by Genscript and cloned to the vector pCMV-IRES-puro. The expression construct was transfected in HEK293F cells using polyethylenimine. The medium was discarded and replaced with FreeStyle 293 medium after 6-8 hours. After incubation in 37 °C with 5.5% CO2 for 5 days, the supernatant was collected and loaded to Ni-NTA resin for purification. The elution was concentrated and further purified by a Superdex 200 column.

### In-Gel Analysis of SARS-CoV-2 S and Human ACE2 Proteins

A 3.5-μg aliquot of SARS-CoV-2 S protein as well as a 2-µg aliquot of human ACE2 were combined with Laemmli sample buffer, analyzed on a 4-12% Invitrogen NuPage Bis-Tris gel using the MES pH 6.5 running buffer, and stained with Coomassie Brilliant Blue G-250.

### Analysis of N-linked and O-linked Glycans Released from SARS-Cov-2 S and Human ACE2 Proteins

Aliquots of approximately 25-50 μg of S or ACE2 protein were processed for glycan analysis as previously described (38,39). For N-linked glycan analysis, the proteins were digested with trypsin. Following trypsinization, glycopeptides were enriched by C18 Sep-Pak and subjected to PNGaseF digestion to release N-linked glycans. Following PNGaseF digestion, released glycans were separated from residual glycosylated peptides bearing O-linked glycans by C18 Sep-Pak. O-glycosylated peptides were eluted from the Sep-Pak and subjected to reductive *β*-elimination to release the O-glycans. Another 25-50 μg aliquot of each protein was denatured with SDS and digested with PNGaseF to remove N-linked glycans. The de-N-glycosylated, intact protein was precipitated with cold ethanol and then subjected to reductive *β*-elimination to release O-glycans. The profiles of O-glycans released from peptides or from intact protein were found to be comparable. N-and O-linked glycans released from glycoproteins were permethylated with methyliodide according to the method of Anumula and Taylor prior to MS analysis (63). Glycan structural analysis was performed using an LTQ-Orbitrap instrument (Orbitrap Discovery, ThermoFisher). Detection and relative quantification of the prevalence of individual glycans was accomplished using the total ion mapping (TIM) and neutral loss scan (NL scan) functionality of the Xcalibur software package version 2.0 (Thermo Fisher Scientific) as previously described (38,39). Mass accuracy and detector response was tuned with a permethylated oligosaccharide standard in positive ion mode. For fragmentation by collision-induced dissociation (CID in MS^2^ and MSn), normalized collision energy of 45% was applied. Most permethylated glycans were identified as singly or doubly charged, sodiated species in positive mode. Sulfated N-glycans were detected as singly or doubly charged, deprotonated species in negative ion mode. Peaks for all charge states were deconvoluted by the charge state and summed for quantification. All spectra were manually interpreted and annotated. The explicit identities of individual monosaccharide residues have been assigned based on known human biosynthetic pathways. Graphical representations of monosaccharide residues are consistent with the Symbol Nomenclature for Glycans (SNFG), which has been broadly adopted by the glycomics community (64). The MS-based glycomics data generated in these analyses and the associated annotations are presented in accordance with the MIRAGE standards and the Athens Guidelines (65). Data annotation and assignment of glycan accession identifiers were facilitated by GRITS Toolbox, GlyTouCan, GNOme, and GlyGen (40,66-68).

### Analysis of Disulfide Bonds for SARS-Cov-2 S Protein by LC-MS

Two 10-μg aliquots of SARS-CoV-2 S protein were denatured by incubating with 20% acetonitrile at room temperature and alkylated by 13.75 mM of iodoacetamide at room temperature in dark. The two aliquots of proteins were then digested respectively using alpha lytic protease, or a combination of trypsin, Lys-C and Glu-C. Following digestion, the proteins were deglycosylated by PNGaseF treatment. The resulting peptides were separated on an Acclaim PepMap RSLC C18 column (75 µm x 15 cm) and eluted into the nano-electrospray ion source of an Orbitrap Fusion(tm) Lumos(tm) Tribrid(tm) mass spectrometer at a flow rate of 200 nL/min. The elution gradient consists of 1-40% acetonitrile in 0.1% formic acid over 370 minutes followed by 10 minutes of 80% acetonitrile in 0.1% formic acid. The spray voltage was set to 2.2 kV and the temperature of the heated capillary was set to 280 °C. Full MS scans were acquired from m/z 200 to 2000 at 60k resolution, and MS/MS scans following electron transfer dissociation (ETD) were collected in the Orbitrap at 15k resolution. The raw spectra were analyzed by Byonic (v3.8.13, Protein Metrics Inc.) with mass tolerance set as 20 ppm for both precursors and fragments. The search output was filtered at 0.1% false discovery rate and 10 ppm mass error. The spectra assigned as cross-linked peptides were manually evaluated for Cys0015 and Cys0136.

### Analysis of Site-Specific N-linked Glycopeptides for SARS-Cov-2 S and Human ACE2 Proteins by LC-MS

Four 3.5-μg aliquots of SARS-CoV-2 S protein were reduced by incubating with 10 mM of dithiothreitol at 56 °C and alkylated by 27.5 mM of iodoacetamide at room temperature in dark. The four aliquots of proteins were then digested respectively using alpha lytic protease, chymotrypsin, a combination of trypsin and Glu-C, or a combination of Glu-C and AspN. Three 10-μg aliquots of ACE2 protein were reduced by incubating with 5 mM of dithiothreitol at 56 °C and alkylated by 13.75 mM of iodoacetamide at room temperature in dark. The three aliquots of proteins were then digested respectively using alpha lytic protease, chymotrypsin, or a combination of trypsin and Lys-C. The resulting peptides were separated on an Acclaim PepMap RSLC C18 column (75 µm x 15 cm) and eluted into the nano-electrospray ion source of an Orbitrap Fusion(tm) Lumos(tm) Tribrid(tm) mass spectrometer at a flow rate of 200 nL/min. The elution gradient consists of 1-40% acetonitrile in 0.1% formic acid over 370 minutes followed by 10 minutes of 80% acetonitrile in 0.1% formic acid. The spray voltage was set to 2.2 kV and the temperature of the heated capillary was set to 280 °C. Full MS scans were acquired from m/z 200 to 2000 at 60k resolution, and MS/MS scans following higher-energy collisional dissociation (HCD) with stepped collision energy (15%, 25%, 35%) were collected in the Orbitrap at 15k resolution. pGlyco v2.2.2 (43) was used for database searches with mass tolerance set as 20 ppm for both precursors and fragments. The database search output was filtered to reach a 1% false discovery rate for glycans and 10% for peptides. Quantitation was performed by calculating spectral counts for each glycan composition at each site. Any N-linked glycan compositions identified by only one spectra were removed from quantitation. N-linked glycan compositions were categorized into 22 classes (including Unoccupied): HexNAc(2)Hex(9∼5)Fuc(0∼1) was classified as M9 to M5 respectively; HexNAc(2)Hex(4∼1)Fuc(0∼1) was classified as M1-M4; HexNAc(3∼6)Hex(5∼9)Fuc(0)NeuAc(0∼1) was classified as Hybrid with HexNAc(3∼6)Hex(5∼9)Fuc(1∼2)NeuAc(0∼1) classified as F-Hybrid; Complex-type glycans are classified based on the number of antenna, fucosylation, and sulfation: HexNAc(3)Hex(3∼4)Fuc(0)NeuAc(0∼1) is assigned as A1 with HexNAc(3)Hex(3∼4)Fuc(1∼2)NeuAc(0∼1) assigned as F-A1; HexNAc(4)Hex(3∼5)Fuc(0)NeuAc(0∼2) is assigned as A2/A1B with HexNAc(4)Hex(3∼5)Fuc(1∼5)NeuAc(0∼2) assigned as F-A2/A1B; HexNAc(5)Hex(3∼6)Fuc(0)NeuAc(0∼3) is assigned as A3/A2B with HexNAc(5)Hex(3∼6)Fuc(1∼3)NeuAc(0∼3) assigned as F-A3/A2B; HexNAc(6)Hex(3∼7)Fuc(0)NeuAc(0∼4) is assigned as A4/A3B with HexNAc(6)Hex(3∼7)Fuc(1∼3)NeuAc(0∼4) assigned as F-A4/A3B; HexNAc(7)Hex(3∼8)Fuc(0)NeuAc(0∼1) is assigned as A5/A4B with HexNAc(7)Hex(3∼8)Fuc(1∼3)NeuAc(0∼1) as F-A5/A4B; HexNAc(8)Hex(3∼9)Fuc(0) is assigned as A6/A5B with HexNAc(8)Hex(3∼9)Fuc(1) assigned as F-A6/A5B; any glycans identified with a sulfate are assigned as Sulfated.

### Analysis of Deglycosylated SARS-Cov-2 S and Human ACE2 Proteins by LC-MS

Three 3.5-μg aliquots of SARS-CoV-2 S protein were reduced by incubating with 10 mM of dithiothreitol at 56 °C and alkylated by 27.5 mM of iodoacetamide at room temperature in dark. The three aliquots were then digested respectively using chymotrypsin, Asp-N, or a combination of trypsin and Glu-C. Two 10-μg aliquots of ACE2 protein were reduced by incubating with 5 mM of dithiothreitol at 56 °C and alkylated by 13.75 mM of iodoacetamide at room temperature in dark. The two aliquots were then digested respectively using chymotrypsin, or a combination of trypsin and Lys-C. Following digestion, the proteins were deglycosylated by Endoglycosidase H followed by PNGaseF treatment in the presence of ^18^O water. The resulting peptides were separated on an Acclaim PepMap RSLC C18 column (75 µm x 15 cm) and eluted into the nano-electrospray ion source of an Orbitrap Fusion(tm) Lumos(tm) Tribrid(tm) mass spectrometer at a flow rate of 200 nL/min. The elution gradient consists of 1-40% acetonitrile in 0.1% formic acid over 370 minutes followed by 10 minutes of 80% acetonitrile in 0.1% formic acid. The spray voltage was set to 2.2 kV and the temperature of the heated capillary was set to 280 °C. Full MS scans were acquired from m/z 200 to 2000 at 60k resolution, and MS/MS scans following collision-induced dissociation (CID) at 38% collision energy were collected in the ion trap. The spectra were analyzed using SEQUEST (Proteome Discoverer 1.4) with mass tolerance set as 20 ppm for precursors and 0.5 Da for fragments. The search output was filtered using ProteoIQ (v2.7) to reach a 1% false discovery rate at protein level and 10% at peptide level. Occupancy of each N-linked glycosylation site was calculated using spectral counts assigned to the ^18^O-Asp-containing (PNGaseF-cleaved) and/or HexNAc-modified (EndoH-cleaved) peptides and their unmodified counterparts.

### Analysis of Site-Specific O-linked Glycopeptides for SARS-Cov-2 S and Human ACE2 Proteins by LC-MS

Three 10-μg aliquots of SARS-CoV-2 S protein and one 10-μg aliquot of ACE2 protein were reduced by incubating with 5 mM of dithiothreitol at 56 °C and alkylated by 13.75 mM of iodoacetamide at room temperature in dark. The four aliquots were then digested respectively using trypsin, Lys-C, Arg-C, or a combination of trypsin and Lys-C. Following digestion, the proteins were deglycosylated by PNGaseF treatment and then digested with O-protease OpeRATOR®. The resulting peptides were separated on an Acclaim PepMap RSLC C18 column (75 µm x 15 cm) and eluted into the nano-electrospray ion source of an Orbitrap Fusion(tm) Lumos(tm) Tribrid(tm) mass spectrometer at a flow rate of 200 nL/min. The elution gradient consists of 1-40% acetonitrile in 0.1% formic acid over 370 minutes followed by 10 minutes of 80% acetonitrile in 0.1% formic acid. The spray voltage was set to 2.2 kV and the temperature of the heated capillary was set to 280 °C. Full MS scans were acquired from m/z 200 to 2000 at 60k resolution, and MS/MS scans following higher-energy collisional dissociation (HCD) with stepped collision energy (15%, 25%, 35%) or electron transfer dissociation (ETD) were collected in the Orbitrap at 15k resolution. The raw spectra were analyzed by Byonic (v3.8.13) with mass tolerance set as 20 ppm for both precursors and fragments. MS/MS filtering was applied to only allow for spectra where the oxonium ions of HexNAc were observed. The search output was filtered at 0.1% false discovery rate and 10 ppm mass error. The spectra assigned as O-linked glycopeptides were manually evaluated. Quantitation was performed by calculating spectral counts for each glycan composition at each site. Any O-linked glycan compositions identified by only one spectra were removed from quantitation. Occupancy of each O-linked glycosylation site was calculated using spectral counts assigned to any glycosylated peptides and their unmodified counterparts from searches without MS/MS filtering.

### Sequence Analysis of SARS-CoV-2 S and Human ACE2 Proteins

The genomes of SARS-CoV as well as bat and pangolin coronavirus sequences reported to be closely related to SARS-CoV-2 were downloaded from NCBI. The S protein sequences from all of those genomes were aligned using EMBOSS needle v6.6.0 (61) via the EMBL-EBI provided web service (69). Manual analysis was performed in the regions containing canonical N-glycosylation sequons (N-X-S/T). For further sequence analysis of SARS-CoV-2 S variants, the genomes of SARS-CoV-2 were downloaded from NCBI and GISAID and further processed using Biopython 1.76 to extract all sequences annotated as “surface glycoprotein” and to remove any incomplete sequence as well as any sequence containing unassigned amino acids. For sequence analysis of human ACE2 variants, the single nucleotide polymorphisms (SNPs) of ACE2 were extracted from the NCBI dbSNP database and filtered for missense mutation entries with a reported minor allele frequency. Manual analysis was performed on both SARS-CoV-2 S and human ACE2 variants to further examine the regions containing canonical N-glycosylation sequons (N-X-S/T). LibreOffice Writer was used to shade regions on the linear sequence of S and ACE2.

### 3D Structural Modeling and Molecular Dynamics Simulation of Glycosylated SARS-CoV-2 S and Human ACE2 Proteins

#### SARS-CoV2 Spike (S) protein structure and ACE2 co-complex

A 3D structure of the prefusion form of the S protein (RefSeq: YP_009724390.1, UniProt: P0DTC2 SPIKE_SARS2), based on a Cryo-EM structure (PDB code 6VSB) (48), was obtained from the SWISS-MODEL server (swissmodel.expasy.org). The model has 95% coverage (residues 27 to 1146) of the S protein. The receptor binding domain (RBD) in the “open” conformation was replaced with the RBD from an ACE2 co-complex (PDB code 6M0J) by grafting residues C336 to V524. *Glycoform generation* – Glycans (detected by glycomics) were selected for installation on glycosylated S and ACE2 sequons (detected by glycoproteomics) based on three sets of criteria designed to reasonably capture different aspects of glycosylation microheterogeneity. We denote the first of these glycoform models as “Abundance.” The glycans selected for installation to generate the Abundance model were chosen because they were identified as the most abundant glycan structure (detected by glycomics) that matched the most abundant glycan composition (detected by glycoproteomics) at each individual site. We denote the second glycoform model as “Oxford Class.” The glycans selected for installation to generate the Oxford Class model were chosen because they were the most abundant glycan structure, (detected by glycomics) that was contained within the most highly represented Oxford classification group (detected by glycoproteomics) at each individual site (**Fig. S7, Supplemental Table, Tabs 1**,**8)**. Finally, we denote the third glycoform model as “Processed.” The glycans selected for installation to generate the Processed model were chosen because they were the most highly trimmed, elaborated, or terminally decorated structure (detected by glycomics) that corresponded to a composition (detected by glycoproteomics) which was present at *≥* 1/3^rd^ of the abundance of the most highly represented composition at each site (**Supplemental Table, Tab 1**). 3D structures of the three glycoforms (Abundance, Oxford Class, Processed) were generated for the SARS-CoV2 S protein alone, and in complex with the glycosylated ACE2 protein. The glycoprotein builder available at GLYCAM-Web (www.glycam.org) was employed together with an in-house program that adjusts the asparagine side chain torsion angles and glycosidic linkages within known low-energy ranges (70) to relieve any atomic overlaps with the core protein, as described previously (71,72).

#### Energy minimization and Molecular dynamics (MD) simulations

Each glycosylated structure was placed in a periodic box of TIP3P water molecules with a 10 Å buffer between the solute and the box edge. Energy minimization of all atoms was performed for 20,000 steps (10,000 steepest decent, followed by 10,000 conjugant gradient) under constant pressure (1 atm) and temperature (300 K) conditions. All MD simulations were performed under nPT conditions with the CUDA implementation of the PMEMD (73,74) simulation code, as present in the Amber14 software suite (University of California, San Diego). The GLYCAM06j force field (75) and Amber14SB force field (76) were employed for the carbohydrate and protein moieties, respectively. A Berendsen barostat with a time constant of 1 ps was employed for pressure regulation, while a Langevin thermostat with a collision frequency of 2 ps^-1^ was employed for temperature regulation. A nonbonded interaction cut-off of 8 Å was employed. Long-range electrostatics were treated with the particle-mesh Ewald (PME) method (77). Covalent bonds involving hydrogen were constrained with the SHAKE algorithm, allowing an integration time step of 2 fs to be employed. The energy minimized coordinates were equilibrated at 300K over 400 ps with restraints on the solute heavy atoms. Each system was then equilibrated with restraints on the C*α* atoms of the protein for 1ns, prior to initiating 4 independent 250 ns production MD simulations with random starting seeds for a total time of 1 μs per system, with no restraints applied.

#### Antigenic surface analysis

A series of 3D structure snapshots of the simulation were taken at 1 ns intervals and analysed in terms of their ability to interact with a spherical probe based on the average size of hypervariable loops present in an antibody complementarity determining region (CDR), as described recently (https://www.biorxiv.org/content/10.1101/2020.04.07.030445v2). The percentage of simulation time each residue was exposed to the AbASA probe was calculated and plotted onto both the 3D structure and primary sequence.

### Analysis of SARS-CoV-2 Spike VSV pseudoparticles (ppVSV-SARS-2-S)

293T cells were transfected with an expression plasmid encoding SARS-CoV-2 Spike (pcDNAintron-SARS-2-SΔ19). To increase cell surface expression, the last 19 amino acids containing the Golgi retention signal were removed. Two SΔ19 constructs were compared, one started with Met1 and the other with Met2. Twenty-four hours following transfection, cells were transduced with ppVSVΔG-VSV-G (particles that were pseudotyped with VSV-G in trans). One hour following transduction cells were extensively washed and media was replaced. Supernatant containing particles were collected 12-24 hour following transduction and cleared through centrifugation. Cleared supernatant was frozen at −80°C for future use. Target cells VeroE6 were seeded in 24-well plates (5×10^5^ cells/mL) at a density of 80% coverage. The following day, ppVSV-SARS-2-S/GFP particles were transduced into target cells for 60 minutes, particles pseudotyped with VSV-G, Lassa virus GP, or no glycoprotein were included as controls. 24 hours following transduction, transduced cells were released from the plate with trypsin, fixed with 4% formaldehyde, and GFP-positive virus-transduced cells were quantified using flow cytometry (Bectin Dickson BD-LSRII). To quantify the ability of various SARS-CoV-2 S mutants to mediate fusion, effector cells (HEK293T) were transiently transfected with the indicated pcDNAintron-SARS-2-S expression vector or measles virus H and F (78). Effector cells were infected with MVA-T7 four hours following transduction to produce the T7 polymerase (79). Target cells naturally expressing the receptor ACE2 (Vero) or ACE2 negative cells (HEK293T) were transfected with pTM1-luciferase, which encodes for firefly luciferase under the control of a T7 promoter (80). 24 hours following transfection, the target cells were lifted and added to the effector cells at a 1:1 ratio. 4 hours following co-cultivation, cells were washed, lysed and luciferase levels were quantified using Promega’s Steady-Glo substrate. To visualize cell-to-cell fusion, Vero cells were co-transfected with pGFP and the pcDNAintron-SARS-2-S constructs. 24 hours following transfection, syncytia was visualized by fluorescence microscopy.

## DATA AVAILABILITY

The mass spectrometry proteomics data are available via ProteomeXchange with identifiers listed in the KEY RESOURCES TABLE.

## SUPPLEMENTAL INFORMATION

Tables (1, 16 tabs), Figures (7), Movies (4), and Simulations (3).

## SUPPLEMENTAL LEGEND

**Supplemental Table Tab 1**. Glycans modeled as Abundance, Oxford Class, and Processed. **Supplemental Table Tab 2**. Cys0015-Cys0136 Disulfide Linked Peptide for SARS-CoV-2 S. **Supplemental Table Tab 3**. Detection of N-linked glycans released from SARS-CoV-2 S and human ACE2. Relative abundance (prevalence) of each species is calculated based on peak intensity in full MS.

**Supplemental Table Tab 4**. Detection of O-linked glycans released from SARS-CoV-2 S and human ACE2. Relative abundance (prevalence) of each species is calculated based on peak intensity in full MS.

**Supplemental Table Tab 5**. N-linked glycan occupancy at each site of SARS-CoV-2 S and human ACE2. Occupancy is calculated using spectral counts assigned to the 18O-Asp-containing (PNGaseF-cleaved) and/or HexNAc-modified (EndoH-cleaved) peptides and their unmodified counterparts. Sequon refers to the Asn-x-Ser/Thr/Cys, Asn-Gly-x sequences.

**Supplemental Table Tab 6**. N-linked glycan compositions identified at each site of SARS-CoV- 2 S and human ACE2. Asn(N)# indicates the numbers of asparagines in protein sequences. In compositions: N=HexNAc, H=Hexose (Hex), F=Fucose (Fuc), A=Neu5Ac, S=Sulfate. In fucosylation: NoFuc=No Fuc identified; 1Core=One Fuc identified at core position; 1Term=One Fuc identified at terminal position; 1Core and 1Term=One Fuc identified as a mixture of core and terminal positions; 1Core1Term=Two Fuc identified and one is at core and the other is at terminal; 2Term=Two Fuc identified at terminal positions; 1Core1Term and 2Term=Two Fuc identified as a mixture of core and terminal positions; 1Core2Term=Three Fuc identified and one is at core and the others are at terminal; 3Term=Three Fuc identified at terminal positions; 1Core2Term and 3Term=Three Fuc identified as a mixture of core and terminal positions; 1Core3Term=Four Fuc identified and one is at core and the others are at terminal; 4Term=Four Fuc identified at terminal positions; 1Core3Term and 4Term=Four Fuc identified as a mixture of core and terminal positions; 1Core4Term=Five Fuc identified and one is at core and the others are at terminal.

**Supplemental Table Tab 7**. N-linked glycan types identified at each site of SARS-CoV-2 S and human ACE2. All N-linked glycans are categorized into 3 types: high-mannose, hybrid and complex.

**Supplemental Table Tab 8**. N-linked glycan oxford classes identified at each site of SARS-CoV-2 S and human ACE2. All N-linked glycan compositions are categorized into 22 classes: M9 to M5 respectively is defined as HexNAc(2)Hex(9∼5)Fuc(0∼1); M1-M4 is defined as HexNAc(2)Hex(4∼1)Fuc(0∼1); Hybrid is defined as HexNAc(3∼6)Hex(5∼9)Fuc(0)NeuAc(0∼1) and F-Hybrid is defined as HexNAc(3∼6)Hex(5∼9)Fuc(1∼2)NeuAc(0∼1). Complex-type glycans are classified based on the number of antenna, fucosylation, and sulfation: HexNAc(3)Hex(3∼4)Fuc(0)NeuAc(0∼1) is assigned as A1 with HexNAc(3)Hex(3∼4)Fuc(1∼2)NeuAc(0∼1) assigned as F-A1; HexNAc(4)Hex(3∼5)Fuc(0)NeuAc(0∼2) is assigned as A2/A1B with HexNAc(4)Hex(3∼5)Fuc(1∼5)NeuAc(0∼2) assigned as F-A2/A1B; HexNAc(5)Hex(3∼6)Fuc(0)NeuAc(0∼3) is assigned as A3/A2B with HexNAc(5)Hex(3∼6)Fuc(1∼3)NeuAc(0∼3) assigned as F-A3/A2B; HexNAc(6)Hex(3∼7)Fuc(0)NeuAc(0∼4) is assigned as A4/A3B with HexNAc(6)Hex(3∼7)Fuc(1∼3)NeuAc(0∼4) assigned as F-A4/A3B; HexNAc(7)Hex(3∼8)Fuc(0)NeuAc(0∼1) is assigned as A5/A4B with HexNAc(7)Hex(3∼8)Fuc(1∼3)NeuAc(0∼1) assigned as F-A5/A4B; HexNAc(8)Hex(3∼9)Fuc(0) is assigned as A6/A5B with HexNAc(8)Hex(3∼9)Fuc(1) assigned as F-A6/A5B; any glycans identified with a sulfate are assigned as Sulfated.

**Supplemental Table Tab 9**. O-linked glycan compositions identified at each site of SARS-CoV- 2 S and human ACE2. Ser/Thr# indicates the numbers of serines or threonines in protein sequences. In compositions: N=HexNAc, H=Hexose (Hex), F=Fucose (Fuc), and A=Neu5Ac.

**Supplemental Table Tab 10**. O-linked glycan occupancy at each site of SARS-CoV-2 S and human ACE2. Occupancy is calculated using spectral counts assigned to the glycosylated peptides and their unmodified counterparts.

**Supplemental Table Tab 11**. SARS-CoV-2 S and human ACE2 variants.

**Supplemental Table Tab 12**. Proteomic Analyses of purified S and ACE2.

**Supplemental Table Tab 13**. Sulfated N-linked glycans released from SARS-CoV-2 S. Following permethylation, almost all of the sulfated hybrid and complex N-glycans are recovered in the organic phase despite their anionic charge. Organic phase permethylated glycans were analyzed by mass spectrometry using negative ion mode. The indicated glycan structures are consistent with the compositions detected at the m/z values shown.

**Supplemental Table Tab 14**. Surface Antigen Exposure of Abundance Glycosylated S. The scale used is 0 (not accessible) to 1.0 (fully accessible).

**Supplemental Table Tab 15**. ACE2-Glycan-S-Peptide Interactions. The scale used is 0 (no interaction) to 1.0 (interacted throughout entire simulation).

**Supplemental Table Tab 16**. S-Glycan-ACE2-Peptide Interactions. The scale used is 0 (no interaction) to 1.0 (interacted throughout entire simulation).

**Supplemental Figure S1**. Defining N-terminus of ACE2 as pyro-glutamine at site Q0018. Representative HCD MS2 spectrum shown.

**Supplemental Figure S2**. Disulfide bond formed between Cysteines 0015 and 0136 of SARS-CoV-2 S. Representative EThcD MS2 spectrum shown.

**Supplemental Figure S3**. Signal P prediction of two different start methionines for SARS-CoV- 2 S.

**Supplemental Figure S4**. Functional characterization of various S constructs in Pseudovirus. (A) Syncytia produced by SARS-CoV-2 S constructs in VeroE6 cells co-transfected with a GFP plasmid to visualize cell-to-cell fusion. Quantification of fusion using a luciferase complementation assay in 293T (B) or VeroE6 cells (C). (D) Transduction efficiency in Vero E6 cells of ppVSV-GFP particles coated in the indicated glycoprotein. Results suggest that start methionine does not alter fusion or efficiency.

**Supplemental Figure S5**. Detection of O-linked glycans released from SARS-CoV-2 S and human ACE2. The detected O-glycans were categorized based on their structures and types. Relative abundance (prevalence) of each species is calculated based on peak intensity in full MS.

**Supplemental Figure S6**. O-linked glycans detected at site T0323 of SARS-CoV-2 S. Representative Step-HCD spectra shown for 6 glycoforms.

**Supplemental Figure S7**. Sequence alignments of SARS-CoV-1 and SARS-CoV-2 S variants as well as alignment of multiple S proteins from related coronaviruses.

**Supplemental Movie A:** Linked to Figure 4C, Glycosylated S antigen accessibility

**Supplemental Movie B:** Linked to Figure 6A, Glycosylated ACE2 with variants

**Supplemental Movie C:** Linked to Figure 7C, Interface of ACE2-S Complex

**Supplemental Movie D:** Linked to Figure 7C, the glycosylated ACE2-S Complex

**Simulation 1:** Linked to Figure 7A, Abundance glycoforms on ACE2-S Complex

**Simulation 2:** Linked to Figure 7A, Oxford class glycoforms on ACE2-S Complex

**Simulation 3:** Linked to Figure 7A, Processed glycoforms on ACE2-S Complex

## Notes

### Competing Interest Statement

The authors have declared no competing interest.

### Summary of Updates

Multiple minor changes including inclusion of sulfated N-glycans, simplification of figures, additional supplemental tables.

