## Supplemental Figures S1-S7 for "Virus-Receptor Interactions of Glycosylated SARS-CoV-2 Spike and Human ACE2 Receptor"

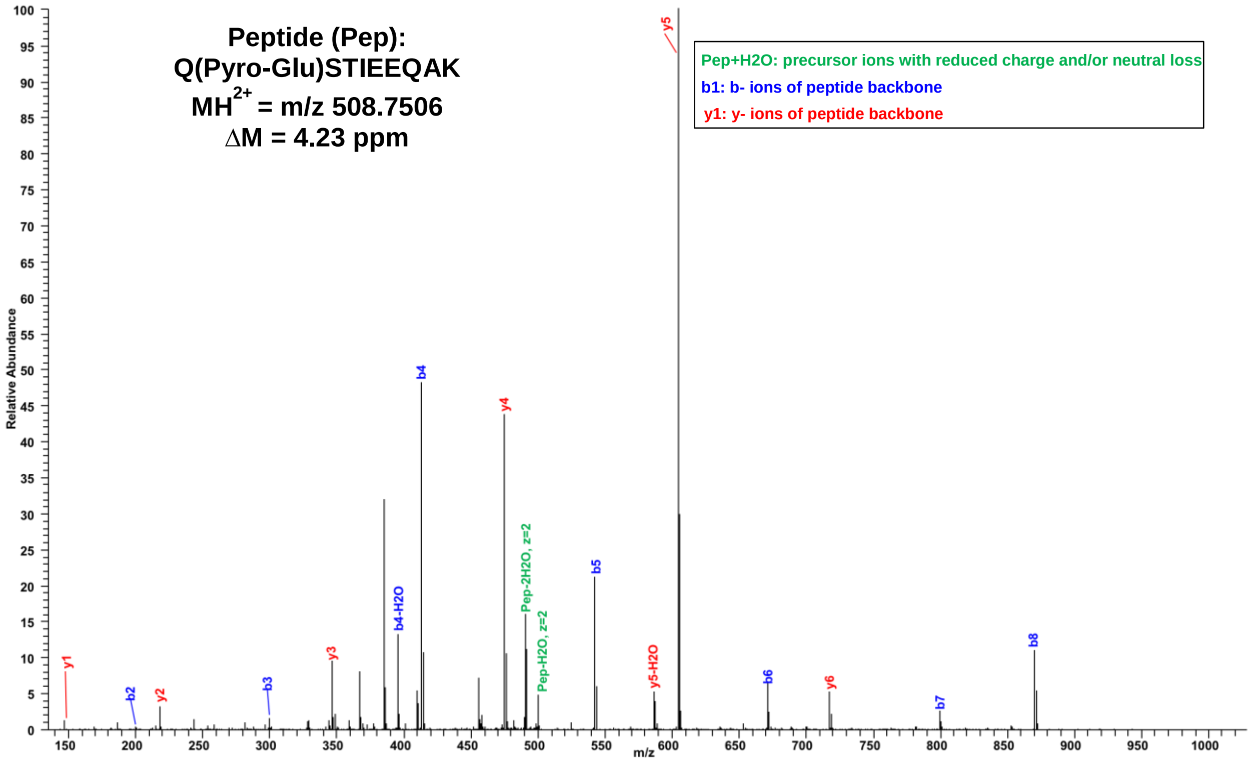
Peptide (Pep):
Q(Pyro-Glu)STIEEQAK
MH2+ = m/z 508.7506
ΔM = 4.23 ppm
Pep+H2O: precursor ions with reduced charge and/or neutral loss
b1: b- ions of peptide backbone
y1: y- ions of peptide backbone
Relative Abundance
m/z
y1
b2
y2
b3
y3
b4-H2O
b4
y4
Pep-2H2O, z=2
Pep-H2O, z=2
b5
y5-H2O
y5
b6
y6
b7
b8

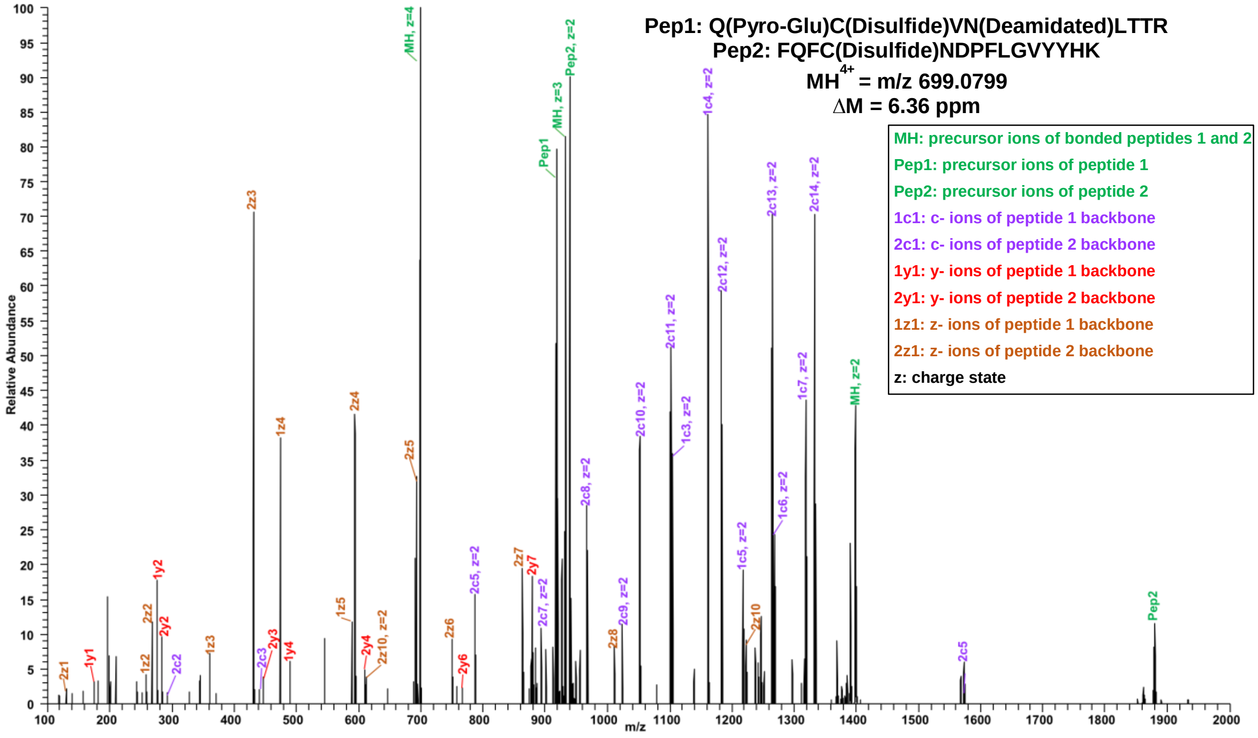
Pep1: Q(Pyro-Glu)C(Disulfide)VN(Deamidated)LTTR
Pep2: FQFC(Disulfide)NDPFLGVYYHK
MH4+ = m/z 699.0799
ΔM = 6.36 ppm
MH: precursor ions of bonded peptides 1 and 2
Pep1: precursor ions of peptide 1
Pep2: precursor ions of peptide 2
1c1: c- ions of peptide 1 backbone
2c1: c- ions of peptide 2 backbone
1y1: y- ions of peptide 1 backbone
2y1: y- ions of peptide 2 backbone
1z1: z- ions of peptide 1 backbone
2z1: z- ions of peptide 2 backbone
z: charge state
Relative Abundance
m/z

- **Utilized Start Met: M**FVFLVLLPLVSSQCVNL…
  - Signal P: SSQC-VN (.54)

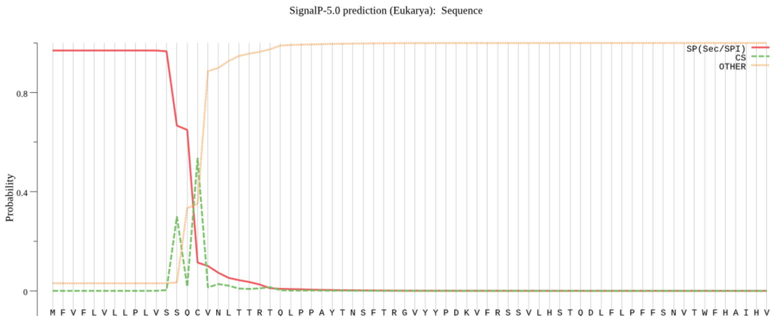


- **Upstream Met**: **M**FLLTTKRT**M**FVFLVLLPLVSSQCVNL…
  - Signal P: SS-QCVN (.64)

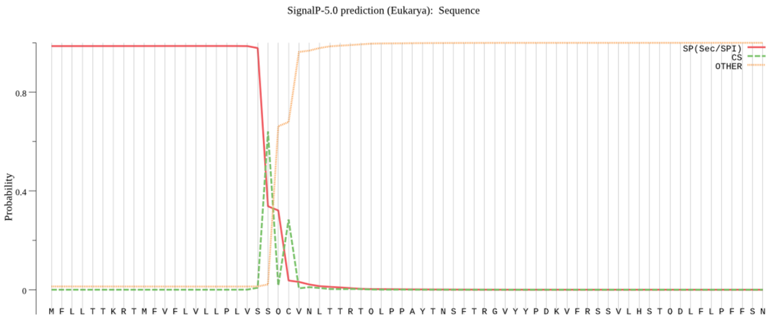

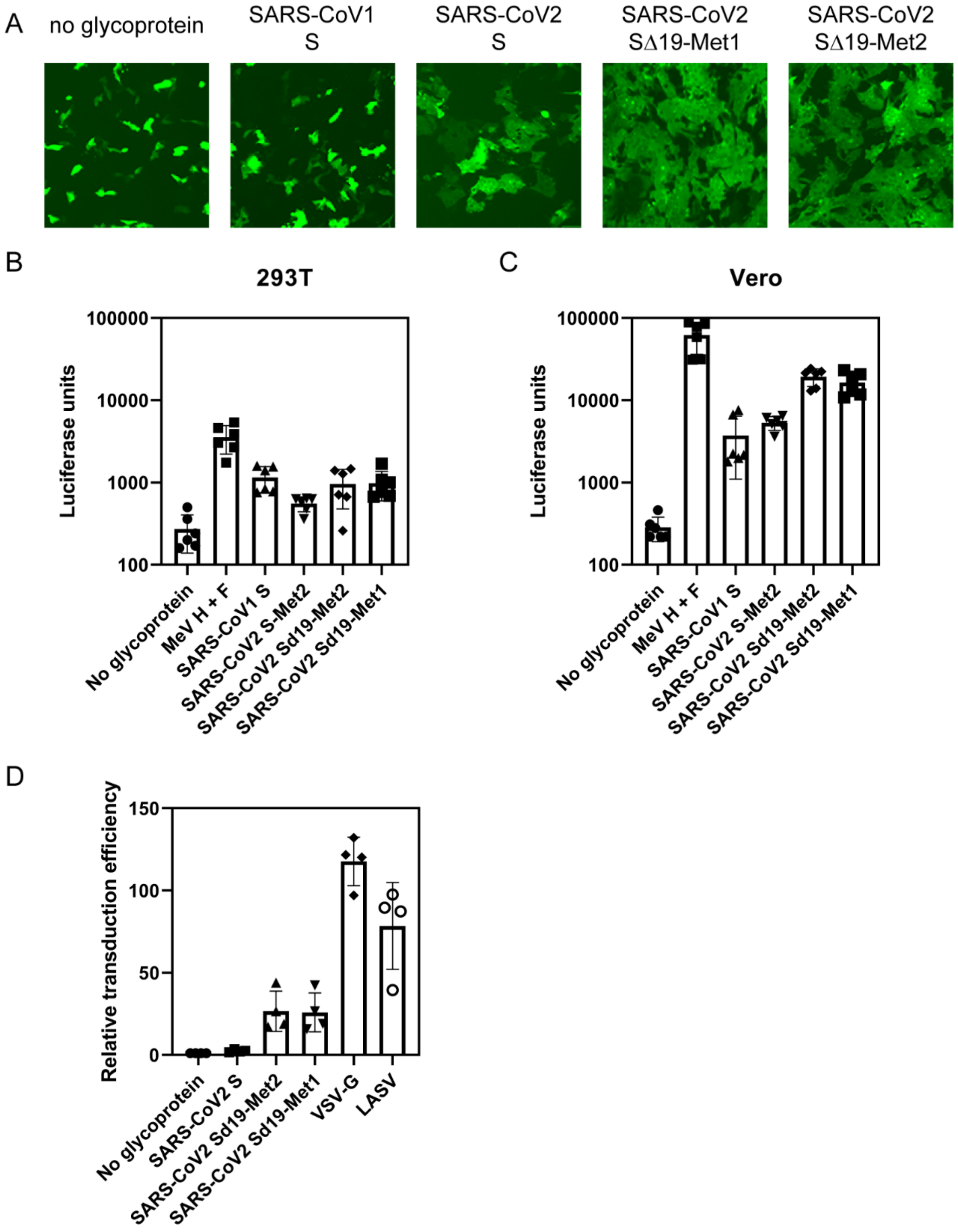
A
no glycoprotein
SARS-CoV1 S
SARS-CoV2 S
SARS-CoV2 SΔ19-Met1
SARS-CoV2 SΔ19-Met2
B
293T
C
Vero
Luciferase units
100000
10000
1000
100
No glycoprotein
MeV H + F
SARS-CoV1 S
SARS-CoV2 S-Met2
SARS-CoV2 Sd19-Met2
SARS-CoV2 Sd19-Met1
D
Relative transduction efficiency
150
100
50
0
No glycoprotein
SARS-CoV2 S
SARS-CoV2 Sd19-Met2
SARS-CoV2 Sd19-Met1
VSV-G
LASV

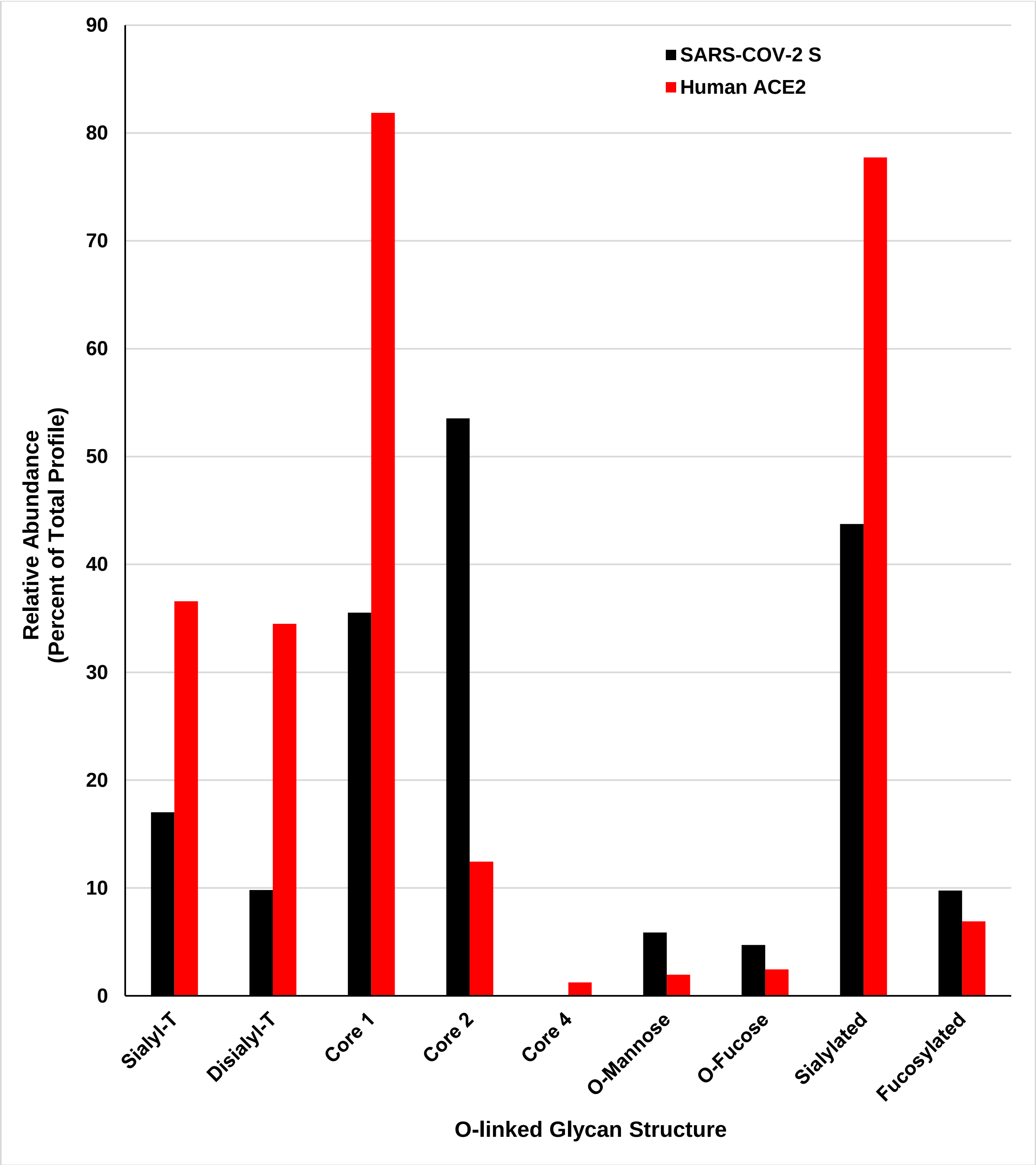

SARS-COV-2 S
Human ACE2
Relative Abundance
(Percent of Total Profile)
0
10
20
30
40
50
60
70
80
90
Sialyl-T
Disialyl-T
Core 1
Core 2
Core 4
O-Mannose
O-Fucose
Sialylated
Fucosylated
O-linked Glycan Structure

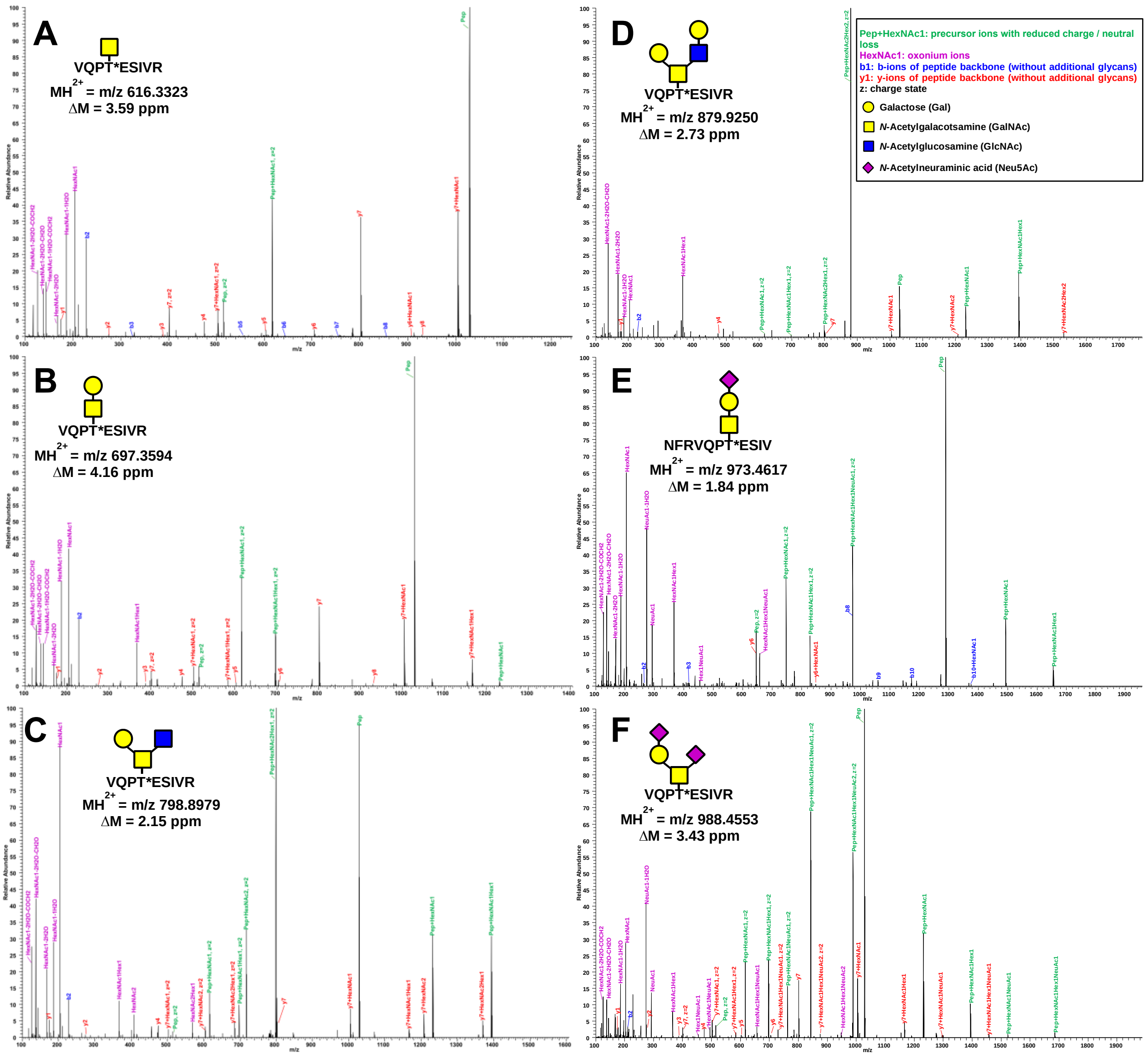
A
VQPT*ESIVR
MH2+ = m/z 616.3323
ΔM = 3.59 ppm
B
VQPT*ESIVR
MH2+ = m/z 697.3594
ΔM = 4.16 ppm
C
VQPT*ESIVR
MH2+ = m/z 798.8979
ΔM = 2.15 ppm
D
VQPT*ESIVR
MH2+ = m/z 879.9250
ΔM = 2.73 ppm
E
NFRVQPT*ESIV
MH2+ = m/z 973.4617
ΔM = 1.84 ppm
F
VQPT*ESIVR
MH2+ = m/z 988.4553
ΔM = 3.43 ppm
Pep+HexNAc1: precursor ions with reduced charge / neutral loss
HexNAc1: oxonium ions
b1: b-ions of peptide backbone (without additional glycans)
y1: y-ions of peptide backbone (without additional glycans)
z: charge state
Galactose (Gal)
N-Acetylgalacotsamine (GalNAc)
N-Acetylglucosamine (GlcNAc)
N-Acetylneuraminic acid (Neu5Ac)
Relative Abundance
m/z

**A**

```
                       ——VAR1——
CoV-1R MFIFLLFLTLTSGSDLDRCTTFDDVQAPNYTQHTSSMRGVYYPDEIFRSDTLYLTQDLFL     60
CoV-2I MFVFLVLLPLVS-SQCVNLTTRTQL-PPAYT--NSFTRGVYYPDKVFRSSVLHSTQDLFL     56
       **:**:.*.*.* *.....**...: .*.**  .*..*******::***..*:.******

CoV-1R PFYSNVTGFHTI-----NHT--FGNPVIPFKDGIYFAATEKSNVVRGWVFGSTMNNKSQS     113
CoV-2I PFFSNVTWFHAIHVSGTNGTKRFDNPVLPFNDGVYFASTEKSNIIRGWIFGTTLDSKTQS     116
       **:****.**.*     *.*  *.***:**.**:***:*****::***:**:*:::*:**
                                         ————————VAR2————————
CoV-1R VIIINNSTNVVIRACNFELCDNPFFAV----SKPMGTQTHTMIFDNAFNCTFEYISDAFS     169
CoV-2I LLIVNNATNVVIKVCEFQFCNDPFLGVYYHKNNKSWMESEFRVYSSANNCTFEYVSQPFL     176
       ::*:**:*****:.*.*:.*::**..*    :.....::...::.:*.******:*..*.

CoV-1R LDVSEKSGNFKHLREFVFKNKDGFLYVYKGYQPIDVVRDLPSGFNTLKPIFKLPLGINIT     229
CoV-2I MDLEGKQGNFKNLREFVFKNIDGYFKIYSKHTPINLVRDLPQGFSALEPLVDLPIGINIT     236
       :*:..*.****:********.**:..:*..:.**::*****.**.:*.:*:..**:*****
         ————VAR3————
CoV-1R NFRAIL----TAFSPAQDI--WGTSAAAYFVGYLKPTTFMLKYDENGTITDAVDCSQNPL     283
CoV-2I RFQTLLALHRSYLTPGDSSSGWTAGAAAYYVGYLQPRTFLLKYNENGTITDAVDCALDPL     296
       .*:.:*    :..:*....  *...****:****:*.**:***:***********:.:**

CoV-1R AELKCSVKSFEIDKGIYQTSNFRVVPSGDVVRFPNITNLCPFGEVFNATKFPSVYAWERK     343
CoV-2I SETKCTLKSFTVEKGIYQTSNFRVQPTESIVRFPNITNLCPFGEVFNATRFASVYAWNRK     356
       :*.**::***.::***********.*:..:*******************:*.*****.**

CoV-1R KISNCVADYSVLYNSTFFSTFKCYGVSATKLNDLCFSNVYADSFVVKGDDVRQIAPGQTG     403
CoV-2I RISNCVADYSVLYNSASFSTFKCYGVSPTKLNDLCFTNVYADSFVIRGDEVRQIAPGQTG     416
       :*************..**********.********:********::**:**********

CoV-1R VIADYNYKLPDDFMGCVLAWNTRNIDATSTGNYNYKYRYLRHGKLRPFERDISNVPFSPD     463
CoV-2I KIADYNYKLPDDFTGCVIAWNSNNLDSKVGGNYNYLYRLFRKSNLKPFERDISTEIYQAG     476
       .************.***:***:.*:*:..*...*****.**..*...*:*******...:...

CoV-1R GKPCT-PPALNCYWPLNDYGFYTTTGIGYQPYRVVVLSFELLNAPATVCGPKLSTDLIKN     522
CoV-2I STPCNGVEGFNCYFPLQSYGFQPTNGVGYQPYRVVVLSFELLHAPATVCGPKKSTNLVKN     536
       ..**. ....***:**..***..*.*:***************:*********.**:*:**

CoV-1R QCVNFNFNGLTGTGVLTPSSKRFQPFQQFGRDVSDFTDSVRDPKTSEILDISPCAFGGVS     582
CoV-2I KCVNFNFNGLTGTGVLTESNKKFLPFQQFGRDIADTTDAVRDPQTLEILDITPCSFGGVS     596
       :***************.*:*:*.********::*.**:****:*.*****:**:*****

CoV-1R VITPGTNASSEVAVLYQDVNCTDVSTAIHADQLTPAWRIYSTGNNVFQTQAGCLIGAEHV     642
CoV-2I VITPGTNTSNQVAVLYQDVNCTEVPVAIHADQLTPTWRVYSTGSNVFQTRAGCLIGAEHV     656
       *******.*::***********:*..*********.**:****:*****:**********
                             VAR4
CoV-1R DTSYECDIPIGAGICASYHTVSLL----R 667
CoV-2I NNSYECDIPIGAGICASYQTQTNSPGGSG 685
       :.***************.*.:..    .
```

**B**

```
AAP13441.1      ... LTSGSDLDRCTTFDDVQA--PNYTQHTSSMRGVYYPD ... FLPFYSNVTGFHTI-----NHT--FGNPVI ...
AGZ48831.1      ... LVSSYTIEKCLDFDDRT--PPANTQFLSSHRGVYYPD ... FLPFDSNVTRFITFGLN-------FDNPII ...
AVP78031.1      ... LVNS----QCVNLTGRTPLNPNYTN--SSQRGVYYPD ... FLPFYSNVSWYYSLTTNNA-ATKRTDNPIL ...
AVP78042.1      ... LVNS----QC-DLTGRTPLNPNYTN--SSQRGVYYPD ... FLPFYSNVSWYYSLTTNNA-ATKRTDNPIL ...
QIA48641.1      ... LVSS----QCVNLTTRTGIPPGYTN--SSTRGVYYPD ... FLPFFSNVTWFNTI--TYQGGSKKFDNPVL ...
QIA48614.1      ... LVSS----QCVNLTTRTGIPPGYTN--SSTRGVYYPD ... FLPFFSNVTWFNTI--NYQGGFKKFDNPVL ...
QIQ54048.1      ... LVSS----QCVNLTTRTGIPPGYTN--SSTRGVYYPD ... FLPFFSNVTWFNTIHLNYQGGFKKFDNPVL ...
QIA48623.1      ... LVSS----QCVNLTTRTGIQPGYTN--SSTRGVYYPD ... FLPFFSNVTWFNTI--NYQGGFKKFDNPVL ...
QIA48632.1      ... LVSS----QCVNLTTRTGIPPGYTN--SSTRGVYYPD ... FLPFFSNVTWFNTI--NYQGGFKKFDNPVL ...
YP_009724390.1  ... LVSS----QCVNLTTRTQLPPAYTN--SFTRGVYYPD ... FLPFFSNVTWFHAIHVSGTNGTKRFDNPVL ...
QHR63300.2      ... LVSS----QCVNLTTRTQLPPAYTN--SSTRGVYYPD ... FLPFFSNVTWFHAIHVSGTNGIKRFDNPVL ...
                    *...    .*  :        *  *:  *  *******     **** ***: : ::           .**::

AAP13441.1      ... FGSTMNNKSQSVIIINNSTNVVIRAC ... PFFAVSKPMGTQTHTM----IFDNAFNCTFEYIS ...
AGZ48831.1      ... FGSTMNNKSQSVIIMNNSTNLVIRAC ... PFF--VVLKSNNTQ-IPSY-IFNNAFNCTFEYVS ...
AVP78031.1      ... FGTTLDNTSQSLLIVNNATNVIIKVC ... PYLSGYYH-NNKTWSIREFAVYSSYANCTFEYVS ...
AVP78042.1      ... FGTTLDNTSQSLLIVNNATNVIIKVC ... PYLSGYYH-NNKTWSIREFAVYSFYANCTFEYVS ...
QIA48641.1      ... FGTTLDARTQSLLIVNNATNVVIKVC ... PFLGVYYHNNNKTWVENEFRVYSSANNCTFEYIS ...
QIA48614.1      ... FGTTLDARTQSLLIVNNATNVVIKVC ... PFLGVYYHNNNKTWVENEFRVYSSANNCTFEYIS ...
QIQ54048.1      ... FGTTLDARTQSLLIVNNATNVVIKVC ... PFLGVYYHNNNKTWVENEFRVYSSANNCTFEYIS ...
QIA48623.1      ... FGTTLDARTQSLLIVNNATNVVIKVC ... PFLGVYYHNNNKTWVENEFRVYSSANNCTFEYIS ...
QIA48632.1      ... FGTTLDARTQSLLIVNNATNVVIKVC ... PFLGVYYHNNNKTWVENEFRVYSSANNCTFEYIS ...
YP_009724390.1  ... FGTTLDSKTQSLLIVNNATNVVIKVC ... PFLGVYYHKNNKSWMESEFRVYSSANNCTFEYVS ...
QHR63300.2      ... FGTTLDSKTQSLLIVNNATNVVIKVC ... PFLGVYYHKNNKSWMESEFRVYSSANNCTFEYVS ...
                    **:*::  :**::*:**:**::*..*     *::       . ::.      ::.   ******:*

AAP13441.1      ... FSLDVSEKSGNFKHLREFVF ... LPLGINITNFRAIL ...
AGZ48831.1      ... FNLDLGEKPGNFKDLREFVF ... LPLGINITNFRTLL ...
AVP78031.1      ... FMLNISGNGGLFNTLREFVF ... LPVSINITKFRTLL ...
AVP78042.1      ... FMLNISGNGGLFNTLREFVF ... LPVSINITKFRTLL ...
QIA48641.1      ... FLMDLEGKQGNFKNLREFVF ... LPIGINITRFQTLL ...
QIA48614.1      ... FLMDLEGKQGNFKNLREFVF ... LPIGINITRFQTLL ...
QIQ54048.1      ... FLMDLEGKQGNFKNLREFVF ... LPIGINITRFQTLL ...
QIA48623.1      ... FLMDLEGKQGNFKNLREFVF ... LPIGINITRFQTLL ...
QIA48632.1      ... FLMDLEGKQGNFKNLREFVF ... LPIGINITRFQTLL ...
YP_009724390.1  ... FLMDLEGKQGNFKNLREFVF ... LPIGINITRFQTLL ...
QHR63300.2      ... FLMDLEGKQGNFKNLREFVF ... LPIGINITRFQTLL ...
                    * :::  : * *: ******     **:.****.*.::* ...
```
